# The trypanosome Variant Surface Glycoprotein mRNA is stabilized by an essential unconventional RNA-binding protein

**DOI:** 10.1101/2020.10.08.331769

**Authors:** Larissa Melo do Nascimento, Franziska Egler, Katharina Arnold, Nina Papavisiliou, Christine Clayton, Esteban Erben

**Affiliations:** Centre for Molecular Biology of Heidelberg University (ZMBH), Im Neuenheimer Feld 280, D69120 Heidelberg, Germany; Division of Immune Diversity, Deutsche Krebsforschungszentrum (DKFZ), Im Neuenheimer Feld 280, D69120 Heidelberg, Germany; Universidad Nacional de San Martín, Campus Miguelete, 25 de Mayo y Francia, San Martín, Provincia de Buenos Aires, Argentina

## Abstract

Salivarian trypanosomes cause human sleeping sickness and economically important livestock diseases. The “bloodstream forms”, which replicate extracellularly in the blood and tissue fluids of mammals, are coated by a monolayer of Variant Surface Glycoprotein (VSG). Switching of the expressed *VSG* gene is central to parasite pathogenicity because it enables the parasites to evade adaptive immunity via antigenic variation. Adequate levels of VSG expression - 10% of total protein and 7% of mRNA - are attained through very active RNA polymerase I transcription, efficient mRNA processing (*trans* splicing of a capped leader and polyadenylation), and high mRNA stability. We here show how *VSG* mRNA stability is maintained. Purification of the *VSG* mRNA with associated proteins specifically selected CFB2, an F-box mRNA-binding protein that lacks known RNA-binding domains. CFB2 binds to a stabilizing complex (MKT1-PBP1-XAC1-LSM12) that recruits poly(A) binding protein and a specialized cap-binding translation initiation complex, EIF4E6-EIF4G5. The interaction of CFB2 with MKT1 is essential for CFB2’s expression-promoting activity, while the F-box auto-regulates CFB2 abundance via interaction with SKP1, a component of the ubiquitination machinery. The results of reporter experiments indicate that CFB2 acts via conserved sequences in the *VSG* mRNA 3’-untranslated region. Depletion of CFB2 leads to highly specific loss of *VSG* mRNA. VSG expression is essential not only for antigenic variation but also for trypanosome cell division. Correspondingly, depletion of CFB2 causes cell cycle arrest, dramatic morphological abnormalities and trypanosome death.

## Introduction

Salivarian trypanosome genomes contain thousands of different *VSG* genes and pseudogenes. The single expressed *VSG* gene is located a telomeric expression site, which consists of an RNA polymerase I promoter, followed by several different “Expression Site Associated Genes” (ESAGs), some repetitive sequences, the *VSG* gene, and finally, telomeric repeats (1–3). Although there are at least ten alternative expression sites, all but one are suppressed by epigenetic mechanisms. Antigenic variation is effected through a combination of expression site transcription switching, and genetic rearrangements that replace the currently expressed *VSG* gene with a different one from a large repertoire of alternative *VSG* genes located at telomeres or in sub-telomeric arrays (4–6). Although different VSGs share their overall structure (7–10), they have relatively limited similarity at the sequence level. The only sequences that are always found in the *VSG* mRNAs of the human and ruminant parasite *Trypanosoma brucei* are an 8mer and a 16mer located in the 3’-untranslated region (3’-UTR). The 16mer sequence is required to maintain the ~2h *VSG* half-life in bloodstream forms (11–13). After uptake into the tsetse fly vector, the trypanosomes convert to the procyclic form; *VSG* transcription is shut off, the mRNA becomes unstable (11,12) and the VSG coat is lost.

## Results

### The *VSG* mRNP proteome

To investigate the mechanism by which *VSG* mRNA is stabilized in bloodstream forms, we purified the *VSG* messenger RiboNucleoProtein complex (mRNP) from cells expressing VSG2. We, like Theil *et al*. (14), adapted methods previously applied to capture proteins associated with the *Xist* noncoding RNA (15–17). After UV crosslinking of proteins to RNA, we used biotinylated 90mers complementary to the entire *VSG* mRNA (Supplementary Table S1) to purify the *VSG* mRNP (Fig. 1A). Alpha tubulin mRNA (*TUB*), which has a similar length and represents about 1.5% of mRNA, was chosen as a control. The two mRNAs were purified sequentially, in alternating order, from the same cell extracts (Fig. 1A) to make background for each as similar as possible; in principle it should be possible to add a third or even a fourth purification to the cascade. At least 1000-fold purification relative to rRNA was obtained (Fig 1B), suggesting final mRNA:rRNA ratios of approximately 1:1 (18). Quantitative mass spectrometry detected 664 different proteins with at least 2 peptides (Supplementary Table S2, sheet 5). 97 proteins were found in all three *VSG* replicates, and 18 in all three *TUB* replicates; this discrepancy probably reflects the relative abundances of the mRNAs in the original lysates. RNA-binding-domain proteins associated with both mRNAs included DRBD18 (19), RBP3 (20), UBP2 (21), DRBD3/PTB1 (22,23) and RBP42 (24). Interestingly, RBP3 was significantly more enriched with *TUB* than with *VSG* (Supplementary Table S2, sheets 3 and 5). RBP3 is essential for normal bloodstream-form trypanosome growth but a previous microarray analysis did not detect the RBP3 - *TUB* mRNA association (20). RBP42, which is a polysome-associated protein that preferentially binds to coding regions (24), also preferred *TUB*. Only one of the two poly(A) binding protein orthologues, PABP2, was reproducibly associated with either *VSG* or *TUB* mRNAs, supporting previously suggested (25) specialized roles for PABP2 and its orthologue PABP1.

**Figure 1.**
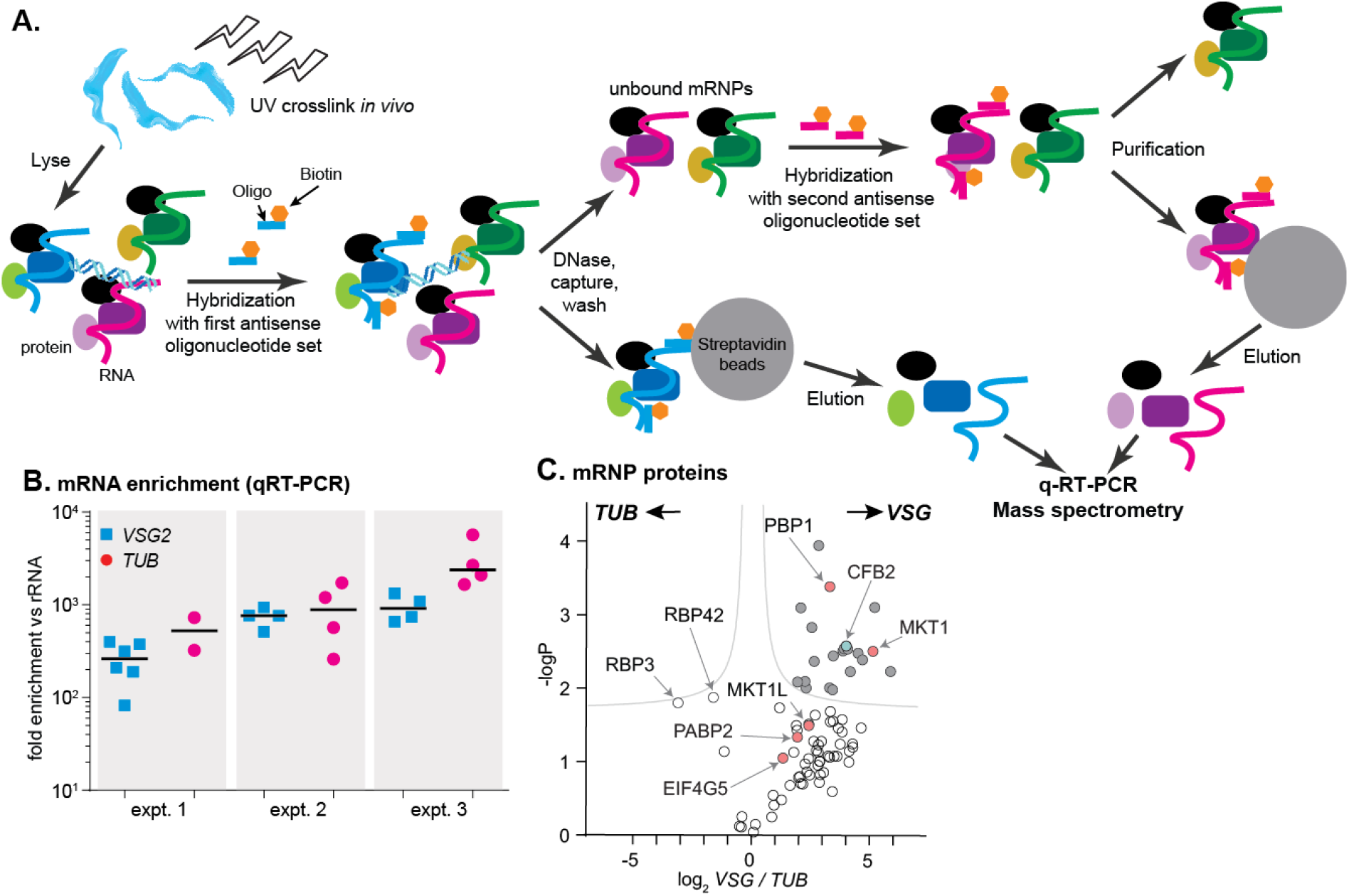
RNA antisense purification identifies proteins that interact directly with VSG mRNA. **A.** A schematic overview of the method. Bloodstream-form T brucei were subjected to UV irradiation. After cell lysis, the lysate was incubated with streptavidin-coated magnetic beads. The unbound portion was then incubated with biotinylated 90 nt long DNA probes, to hybridize to either alpha-tubulin (TUB) or VSG2 mRNA, then probe-target complexes were captured by streptavidin-coated magnetic beads. To decrease background, samples were treated with DNase I. The supernatant was collected, and the second set of ribonucleoprotein complexes (VSG2 or TUB) was captured in a similar way. Protein and RNA were eluted from the beads, and subjected to mass spectrometry (LC-MS/MS) for protein identification and RT-qPCR for relative RNA quantification. **B.** Enrichment of VSG2 and TUB transcripts after RNA antisense purification. The individual data points show RNA levels relative to rRNA for each independent pulldown, measured by RT-qPCR, and bars represent means. Experiment 1 includes some preparations that were purified using only VSG. Before mass spectrometry of VSG or TUB preparations from each experiment, the material for all pull-downs shown was pooled. **C.** The triplicate experiments identified proteins that reproducibly enriched with VSG2 relative to TUB, and vice-versa. Proteins significantly enriched (FDR 1%; s0=0.1) are filled dark grey. Proteins associated with the MKT1 complex are in pink and CFB2 is cyan. The data for this graph are in Supplementary Table S2, sheet 2.

Up to 43 proteins were significantly enriched in the *VSG2* mRNP, depending on the criteria applied (Supplementary Table S2, sheet 1, Fig. 1C). Of these, all but three were detected in the total poly(A)+ mRNP proteome (26), supporting direct binding to mRNA *in vivo*. The proteins with general mRNA-related functions - PABP2, helicases, translation factors and ribosomal proteins - were probably enriched because *VSG* mRNA is more abundant than *TUB. VSG* mRNA-associated proteins with known RNA-binding domains were ALBA3, ZC3H28, ZC3H32, ZC3H41. ALBA3 has previously been implicated in translation enhancement and developmental regulation in procyclic forms of the parasite (27,28), and ZC3H41 as associated with SLRNA, the precursor for mRNA *trans* splicing (29). Although ZC3H32 is bloodstreamform-specific and essential, a tagged version showed no evidence of specificity in mRNA binding (30). The roles of ZC3H28 and ZC3H41 are not known.

To find candidates for stabilization of *VSG* mRNA, we focused on proteins that were highly enriched in the *VSG* mRNP, are known to bind well to mRNA, and are expressed only in the bloodstream form. We also chose proteins that are capable of increasing mRNA stability or translation when artificially “tethered” to an mRNA. In this assay, we express the protein of interest as a fusion with the lambdaN peptide, using cells that also express a reporter mRNA that contains 5 copies of the boxB sequence, which binds the lambdaN peptide with very high affinity. One possible candidate was ERBP1, but this shows only moderate developmental regulation; its co-purification with the *VSG* mRNP may be linked to its association with the endoplasmic reticulum (31). The remaining candidate was CFB2 (cyclin F-box protein 2, Tb927.1.4650).

### CFB2 is associated with *VSG* mRNA

The *CFB2* gene is downstream of several genes encoding CFB1, a related protein (32). Both *CFB1* and *CFB2* mRNAs are much more abundant in bloodstream forms than in procyclic forms, but mass spectrometry (33) and ribosome profiling results (34,35) suggest that CFB2 predominates in bloodstream forms. *CFB2* mRNA persists in stumpy-form trypanosomes (36,37), which are growth-arrested but VSG-expressing bloodstream forms that are poised for differentiation to the procyclic form. Within the Tsetse fly, *CFB2* mRNA is present only in forms that express VSG (38–40), whereas *CFB1* mRNA is also present in the epimastigote form, which lacks VSG (40). *CFB* genes are Kinetoplastid-specific; at least one *CFB* gene is present in all *Trypanosoma* and in *Paratrypanosoma*, but they are absent in *Leishmania*. Alignments of the different regions of the protein suggest that different copies, where present, arose by duplication after species divergence (Supplementary Fig 1A). It is notable that all Salivaria have several *CFB* genes whereas the intracellular *Stercoraria*, which lack antigenic variation, have only one. Moreover, it was already known that depletion of CFB2 from bloodstream forms caused rapid G2 arrest (32) - a phenotype that was also seen after RNAi targeting *VSG* mRNA (41).

To confirm association of CFB2 with *VSG* mRNA we integrated boxB loops into the actively expressed *VSG* gene immediately after the *VSG2* stop codon, preserving the endogenous *VSG* 3’-UTR and polyadenylation site (Fig. 2A). We co-expressed a chimeric GFP protein bearing a streptavidin binding peptide at the C-terminus and, at the N-terminus, the lambdaN peptide (N-GFP-SBP) (Fig. 2A, B); expression was tetracycline-inducible. In addition, 6x-myc-CFB2 was constitutively expressed in the same cells. Affinity purification on a streptavidin matrix allowed us to pull down the *VSG-boxB* mRNA using N-GFP-SBP. Detection of Myc-CFB2 in the pull-down depended on the presence of both lambda-GFP-SBP (Fig 2B), and the *VSG2*-associated *boxBs* (Fig 2C, Supplementary Fig 2A). Other abundant proteins such as RBP10 (42,43) and aldolase (44) were not enriched (Fig 2B,C). In a converse experiment, we used cells expressing CFB2-6xmyc but no boxB reporter. After immunoprecipitation, *VSG2* mRNA was enriched 1.8-3.8 fold relative to rRNA in 4/5 experiments (Supplementary Fig 2B).

**Figure 2.**
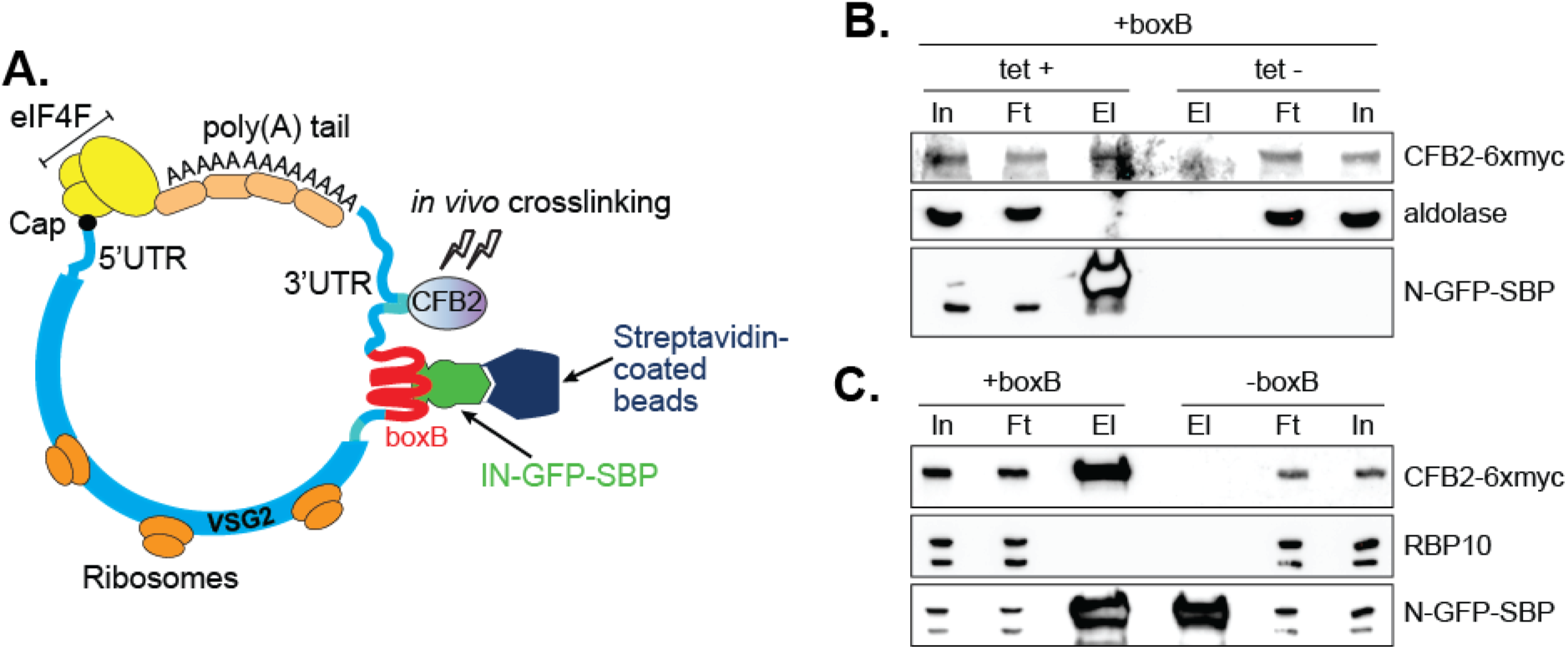
CFB2 associates with VSG2 mRNA. **A.** Method used. The lambda-GFP-SBP binds to boxB aptamer-tagged *VSG2* mRNAs via the lambda moiety located at the N-terminal and interacts with immobilized streptavidin via the SBP moiety at the C-terminus. **B.** CFB2-myc is pulled down with boxB-containing *VSG2* mRNA. Bloodstream cells expressing boxB-tagged *VSG2* mRNA from the active expression site, CFB2-6xmyc from the *RRNA* locus and tetracycline-inducible lambdaN-GFP-SBP were grown; one culture was treated with tetracycline for 6h (+tet) and the other was not (-tet). The lambdaN-GFP-SBP, with associated mRNA and protein, was purified from cell lysates (25 mg of the total protein). For Western analysis, 85% of the eluates (El; corresponding to ~21 mg of input protein) and 40 μg total input (In) or flowthrough (Ft) proteins were resolved by SDS-PAGE and analysed by Western blotting using specific antibodies (Eluate: flowthrough loading of 500:1). Panel B shows that pull-down of CFB2-6xmyc was dependent on the presence of lambdaN-GFP-SBP. Raw data for this Figure are in Supplementary Fig 2A. **C.** This is similar to B, except that all cells expressed lambdaN-GFP-SBP, but one line did not have boxB sequences in the *VSG* mRNA. This shows that pull-down of CFB2-6xmyc by lamb-daN-GFP-SBP depended on the boxB sequences.

### The MKT1-PBP1 complex is associated with *VSG* mRNA

CFB1 and CFB2 have three possible functional domains: the cyclin F-box; a conserved central region; and the MKT1 interaction motif (Fig 3A). Cyclin F-box proteins interact with SKP1 and form an E3 ubiquitin ligase. The residues required for the interaction in Opisthokonts (45–47) are conserved in CFB2 (Fig 3B). The interaction of CFB2 with *T. brucei* SKP1 was confirmed in a yeast 2-hybrid assay (Fig 3C), and was prevented by mutation of critical F-box residues (Supplementary Fig 3).

**Figure 3.**
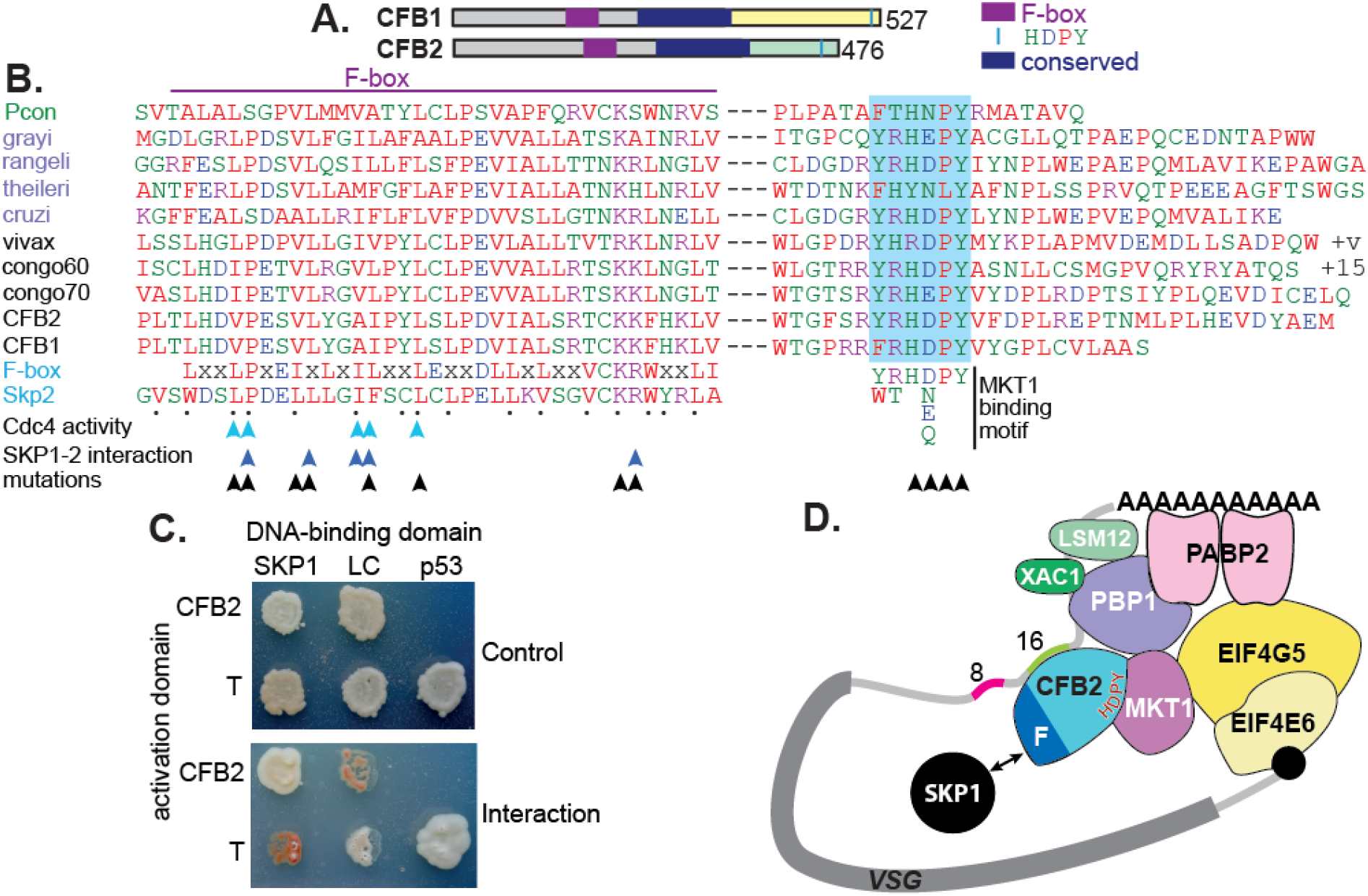
CFB2 has three conserved domains. **A.** Schematic depiction of T brucei CFB1 and CFB2. Similar sequences outside the F-box and HDPY region are in grey and regions conserved in Trypanosoma and Paratrypanosoma are in other colors as indicated. **B.** Sequence alignments created in ClustalOmega. The F-box is on the left and the region with C-terminal HNPY (highlighted in blue) on the right. The colour code for residues is red: nonpolar; green: polar; blue: acidic; purple: basic. The Paratrypanosoma sequence (green label) is at the top. Stercoraria (*T grayi, T. rangeli, T. theileri* and *T. cruzi*) are labelled in mauve and Salivaria are in black (*T vivax, T. congolense*, and *T. brucei* CFB1 and CFB2). Most organisms shown have several very similar CFB gene copies; only one is shown unless sequences are different. “+v” indicates variable C-termini of different *T. vivax* orthologues; “+15” indicates 15 more amino acid residues. At the bottom are a mammalian F-box consensus (“F-box”) and human SKP2. The points below SKP2 indicate residues implicated in its interaction with SKP1 (46). Cyan arrows indicate residues required for yeast Cdc4 activity (45) and dark blue arrows the residues required for the human SKP1-SKP2 interaction (47). Black arrows indicate residues mutated to alanine in our studies. Phosphorylation of CFB2 was detected at tyrosine 431, 16 residues N-terminal to the HDPY signal (52). **C.** The F-box of CFB2 interacts with SKP1: yeast 2-hybrid assay. Fusions of SKP1, LaminC (negative control) and p53 with the DNA-binding domain were co-expressed with CFB2 or SV40 T-antigen fused with the transcription activation domain. The interaction between SV40 T-antigen and p53 is the positive control. The upper panel shows selection of transformants on “double drop-out” medium and the lower panel shows selection for the interaction on “quadruple drop-out” medium. **D.** A model for CFB2 function. The HDPY motif of CFB2 interacts with MKT1, which recruits PBP1 and EIF4G5. PBP1 recruits PABP2, XAC1 and LSM12: and EIF4G5 is complexed with EIF4E6. Meanwhile, the F-box of CFB2 interacts with SKP1; this might, or might not, be compatible with MKT1 and mRNA binding.

The C-terminus of CFB2 includes a motif, RYRHDPY, which is required for the interaction of CFB2 with MKT1 (48,49) (Supplementary Fig 3). *T. brucei* MKT1 forms a complex with PBP1, LSM12, XAC1 and PABP2 (48,49), and also preferentially recruits one of the six alternative cap-binding translation initiator factor complexes, EIF4E6-EIF4G5 (49,50) (Fig 3D). Recruitment of the MKT complex stabilizes bound mRNAs and promotes translation (48). Evidence so far suggests that although both MKT1 and PBP1 have some intrinsic RNA-binding activity, they are recruited to specific mRNAs by various different RNA-binding proteins, resulting in enhanced mRNA abundance and translation (26,48,51). CFB2 showed clear-co-purification with MKT1 and XAC1 (48,49). Correspondingly, both MKT1 and PBP1 were highly enriched in the *VSG* mRNP (Fig. 1C, Supplementary Table S2, sheet 1). EIF4G5, EIF4E6, and EIF4G4 were also detected exclusively with VSG, but the cap-binding partner of EIF4G4, EIF4E3, was not detected. We therefore hypothesized that recruitment of CFB2 to the *VSG* mRNA results in cooperative assembly of an mRNP that includes the MKT complex, PABP2 and EIF4E6-EIF4G5 (Fig. 3D).

### Depletion of CFB2 causes selective loss of *VSG* mRNA

To test our hypothesis, we first examined the effects of CFB2 depletion. We had previously shown that RNAi-mediated depletion of CFB2 resulted in almost immediate G2 arrest, with an accumulation of flagella and basal bodies (32). To find out whether this was accompanied by a specific reduction in *VSG* mRNA, we induced *CFB2* RNAi and measured mRNAs by realtime PCR 3h and 6h later. Tubulin mRNA was unaffected but *VSG* mRNA was already reduced within 3h of *CFB2* RNAi induction (Fig 4A, B). Importantly, preliminary data showed that this was also true for a cell line expressing *VSG3* followed by the 3’-UTR of *VSG2* (Supplementary Fig 2C,D), and transient transfection of *CFB2* double-stranded RNA into cells expressing VSG222 also caused *VSG222* mRNA reduction (Supplementary Fig 2E), suggesting that the *CFB2* RNAi effect does not require a specific *VSG* coding region. Inhibition of VSG synthesis is known to induce translation arrest within 24h (53), but no general translation inhibition was observed over the first 8h of *CFB2* RNAi (Supplementary Fig 2F).

**Figure 4.**
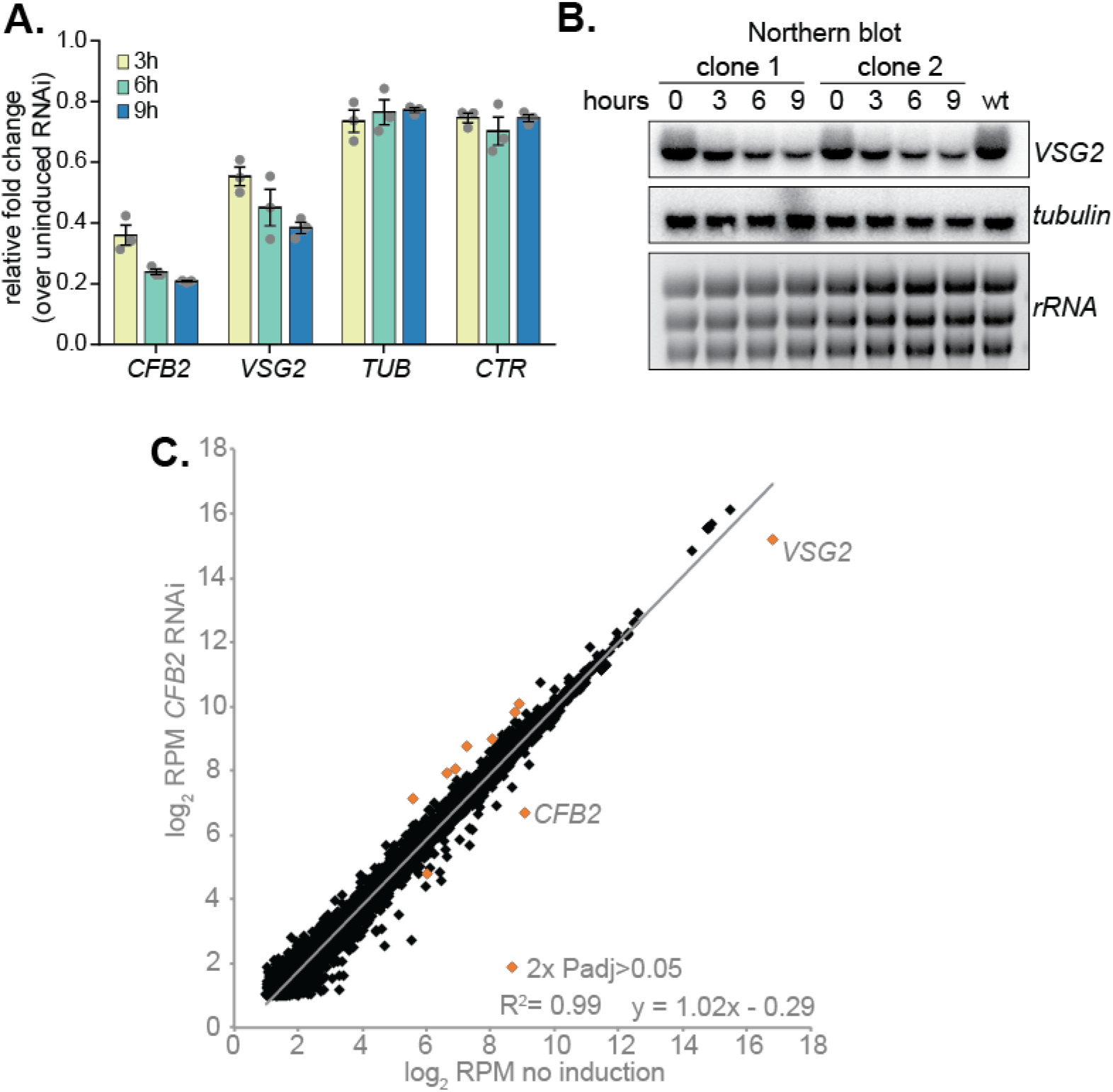
Depletion of CFB2 results in selective loss of VSG mRNA. **A.** Relative quantification of RNA transcript levels corresponding to the active *VSG2* and alphatubulin (*TUB*) mRNAs detected at different sampling time-points (3, 6, 9h) after induction of CFB2 RNAi. CTR (co-transposed region) is a sequence located upstream of the VSG gene, which is transcribed from the same promoter but present only in the mRNA precursor. Results were derived from three biological replicate experiments with standard deviation indicated with error bars, normalised against 18S. **B.** Representative Northern blot analysis of transcript levels after depletion of CFB2 in cells expressing VSG2; details as in (A). Raw data for this Figure are in Supplementary Fig 2G. **C.** Effect of *CFB2* RNAi on the transcriptome. Cells with tetracycline-inducible *CFB2* RNAi were used. The average reads per million reads from two replicates were plotted for each open reading frame. Values below 2 were excluded. Results for Lister427 coding regions are shown (Supplementary Table S3). Data for genes which gave an RPM ratio of more than 2-fold increase or decrease, and a DeSeq2 adjusted P-value of less than 0.05, are highlighted in orange.

We next examined the effect of CFB2 reduction on the transcriptome. After 9h of CFB2 depletion, the only mRNAs that were significantly reduced were those encoding CFB2, VSG2 and, to a slightly lesser extent, Tb927.8.1945 (Fig 4E, Supplementary Tables S3 and S4, Supplementary Fig 4). As previously shown (32) *CFB2* but not *CFB1* transcripts decreased after RNAi induction; the minor decrease in coverage over the *CFB1* genes can be assigned to sequences that are also present in *CFB2* (Fig. 5A). Fig 5B shows reads over the active VSG expression site: the active *VSG2* mRNA was decreased, while mRNAs from the co-transcribed *ESAG* genes were unaffected. This indicates that the effect on *VSG2* mRNA most likely operates at the post-transcriptional level, although we cannot rule out the possibility that some *ESAG* reads come from copies elsewhere in the genome. The level of the *VSG* pre-mRNA, as judged by the *VSG2* co-transposed region (CTR) measurement, also remained constant (Fig 4A).

**Figure 5.**
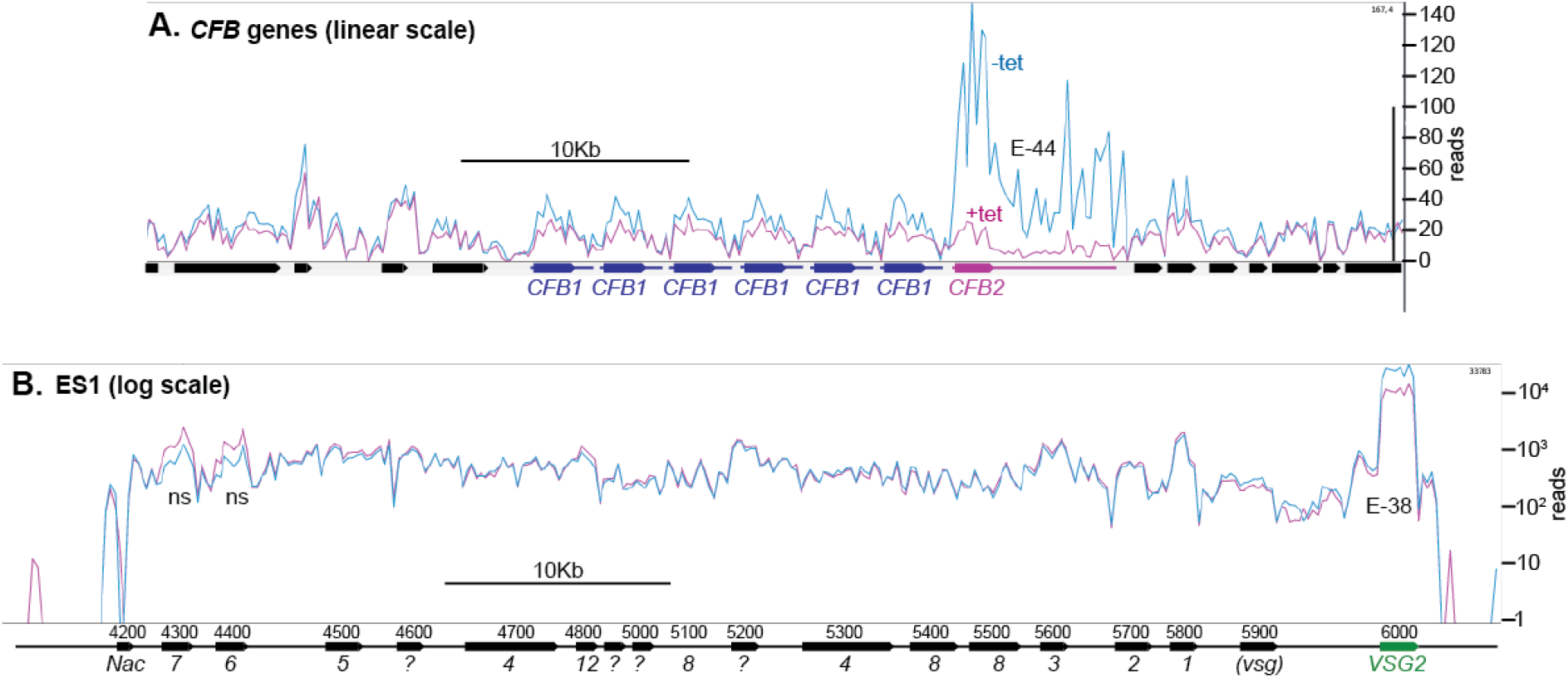
Depletion of CFB2 does not affect expression site transcripts. Reads were aligned to the Lister427 genome, allowing one match per read, and visualised using Artemis. Reads from two samples, one with, and one without, tetracycline, were compared because they had similar overall read counts. **A.** *CFB1* and *CFB2* genes, shown on a linear scale. CFB1 open reading frames are in blue and CFB2 in magenta, and the approximate extents of the untranslated regions are indicated by lines. *CFB1* and *CFB2* share some coding region sequence, but the untranslated regions are different. After tetracycline addition, some reads seen over the *CFB2* open reading frame probably actually originate from *CFB1* genes. E values are the significance of the change from DeSeq2. **B.** Reads aligned over bloodstream-form expression site 1 (BES1), shown on a log10 scale. Most *ESAG* genes are present in multiple copies in the genome, in expression sites and elsewhere. *ESAG* mRNAs are less abundant and presumably less stable than the *VSG* mRNA. ESAGs are normally transcribed from the expression site, although some matching mRNAs may arise from copies elsewhere in the genome. The effect of *CFB2* RNAi was exclusive to VSG2.

Interestingly, thirteen different mRNAs were >1.5x increased after CFB2 depletion (Fig 4E, Supplementary Table 4). Their products included ESAG5-related proteins, cysteine peptidases, the surface protease MSPC (54) and the invariant surface glycoprotein ISG65 (55,56). Some other possible membrane protein mRNAs (57) were also slightly (>1.3x), but significantly, increased. We speculate that loss of *VSG* mRNAs might allow mRNAs encoding other membrane proteins increased access to the secretory pathway; this might facilitate translation, which in turn might indirectly enhance mRNA stability. However, for the remaining increased mRNAs there is no evidence for association with the secretory pathway (58).

### Depletion of CFB2 causes accumulation of cells containing internal flagella

Depletion of *VSG* mRNA by RNAi causes not only a G2 block but also, after 60h after RNAi induction, multiple internal flagella (41). The effects of *CFB2* RNAi were similar, but much more rapid. Cells started to accumulate at G2/M almost immediately, with cell death commencing after about 24h (32). The effects of *CFB2* RNAi are presumably fast because there are only about 5 *CFB2* mRNAs per cell (59) and the protein is unstable (see below). 16h after RNAi induction, the cells had numerous flagella and basal bodies (32). Electron microscopy revealed that at this time, nearly all cells had several external flagella, including both an axoneme and a paraflagellar rod. In most cases there were also flagella inside the cells; sometimes these had enclosing membranes, but often, they did not-exactly the same as was seen depletion of *VSG* mRNA (Fig 6, Supplementary Figs 5 & 6). The trypanosome flagellum emerges into an enclosed structure called the flagellar pocket, which is the site all exocytosis and endocytosis (60). CFB2-depleted cells often had grossly enlarged flagellar pockets. This defect was previously observed after depletion of clathrin or actin, when it is caused by a failure in endocytosis (61,62), and also in cells with flagellar assembly defects (63).

**Figure 6.**
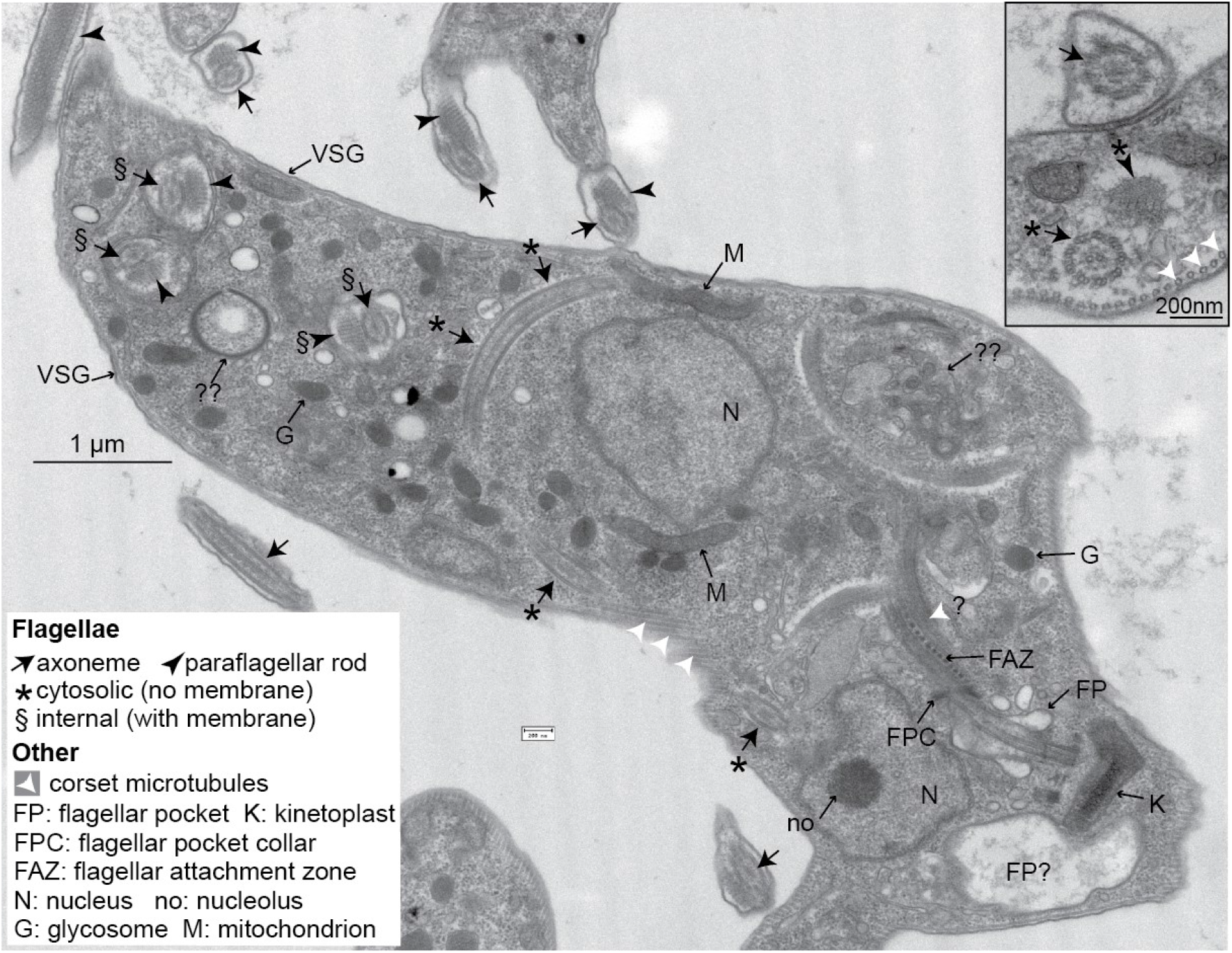
Trypanosomes lacking CFB2 have internal flagella. **A.** A typical trypanosome 16h after induction of RNAi targeting *CFB2*. The central cell has one flagellar pocket of normal appearance (FP); the base of the emerging flagellum can be seen. This should exit the cell at the flagellar pocket collar (FPC), and may then run beneath the indicated flagellar attachment zone (FAZ). Next to the neighbouring kinetoplast (K), which has normal morphology, there is a second vacuole that contains low-density material; this could be part of a second, much enlarged, flagellar pocket. (The usual flagellar pocket diameter does not exceed 500 nm (64)). The cell has two nuclei (N), and in one the section passes through the nucleolus (no). The cell contains at least three additional flagellar axonemes. One (*) is found in both longitudinal as well as transverse cross-section, and lacks a surrounding membrane. Additional axonemes with paraflagellar rods (§) are surrounded by double membranes that may lack the thick VSG coat (indicated at two positions on the cell exterior). The paraflagellar rod is normally seen only on flagella that have exited the flagellar pocket. The glycosomes (G) and mitochondrion (M) appear normal. Double question marks indicate abnormal membrane structures that are suggestive of autophagy and internal membrane proliferation. The inset shows part of an enlarged flagellar pocket with a membrane-enclosed axoneme; beneath it are the four specialised microtubules that are found next to the FAZ; these usually, however, are associated with endoplasmic reticulum. The section also includes a membraneless internal flagellum and flagellar rod. More images are in Supplementary Figures S5 and S6.

### The F-box is implicated in auto-regulation and MKT interaction is required for mRNA activation

Results of a high-throughput study suggested that the half-life of untagged CFB2 is probably less than 1h (65), so we wondered whether the interaction of CFB2 with SKP1 provokes its own degradation. An affinity-purified antibody faintly recognized a ~50 KDa protein in Western blots of trypanosome lysates, probably with less than thousand molecules per cell (Supplementary Figure 7A, B). The abundance of this protein was increased by prior incubation of the cells with MG132 (Supplementary Figure 7C), suggesting that it is unstable. Tagging of both the N and C-terminus did not increase the abundance (Supplementary Figure 7D) or the effect of MG132 Supplementary Figure 7E). We therefore suspected that the F-box interaction with SKP1 might be causing CFB2 instability.

Results from tethering screens had suggested that the C-terminal portion of CFB2 is able to activate expression of a reporter mRNA but the full-length protein does not (26,66). All fragments with activation function contained the C-terminal MKT1 interaction domain, YRHDPY, and lacked the F-box (Supplementary Fig 8A); but many also included a conserved region with basic and hydrophobic residues (illustrated in Supplementary Fig 8B). To examine the functions of both the SKP1 and MKT1 interaction motifs, we expressed various mutants as fusion proteins, with the lambdaN peptide at the N-terminus and a myc tag at the C-terminus, testing their expression and ability to enhance expression of a boxB reporter. Both deletion and mutation of the F-box motif increased fusion protein abundance (Figure 7B, Supplementary Fig 7D and Supplementary Fig 9), implicating the F-box in autoregulation of the stability ectopically expressed CFB2 protein. Increased abundance may explain why the full-length F-box mutant was able to activate reporter expression better than the wild-type protein, although not as much as the C-terminal fragment (Figure 7C, Supplementary Fig 9D). As expected, an intact MKT1 interaction motif was essential for activation. The lack of activation by the full-length protein could at least in part be caused by recruitment of SKP1 to the tethered, RNA-bound protein: we do not know whether this activity is prevented when CFB2 is bound via its own RNA-binding domain.

**Figure 7.**
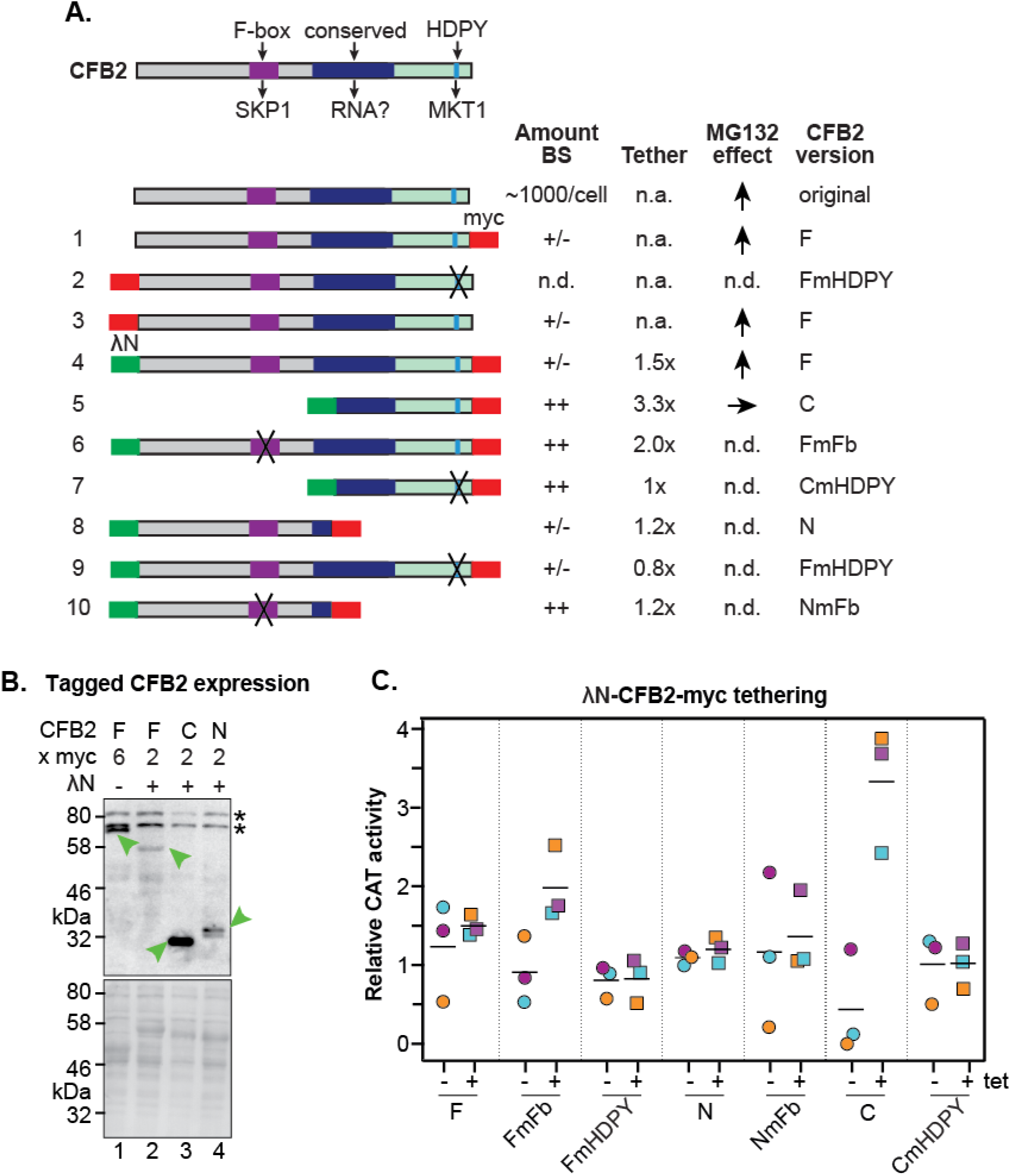
Roles of the SKP1 and MKT1 interaction domains of CFB2. **A.** Structures of different expressed versions of CFB2, with expression level, effects when tethered, and effect of MG132 on the expression level as shown in panels B and C. For expression, +/- means difficult to detect, and ++ is very easy to detect using anti-myc antisera, with double myc tags at either N- or C-terminus as indicated. Constructs 1 and 3 exist in two versions. Versions with an rRNA promoter and 6xmyc tags were used for measurement of the MG132 effect in bloodstream forms. Versions with an inducible promoter and two myc tags were used to assess the effect of CFB2 expression in procyclic forms. All other constructs have two myc tags and an inducible promoter. Versions 4-10 have the N-terminal lambdaN peptide. “F” means “full length”, “C” means C-terminal portion, “N” means N-terminal portion. Mutations in the F-box (Fb) and in the HDPY motif are illustrated in Figure 3. **B.** Expression of fusion proteins in bloodstream forms. Lane 1 shows constitutive expression of CFB2-6xmyc (construct 1) from an rRNA promoter, while lanes 2-4 show expression of different lambdaN-2xmyc-tagged fragments (4, 5, 8) after 24h induction of expression from a tetracycline-inducible EP procyclin promoter. The myc epitope was detected (green arrow). Whole-cell proteins were visualised by Ponceau (bottom). **C.** Graph showing effects of tethering different versions of CFB2 (constructs 4-10). The bloodstream-form trypanosomes used expressed a *CAT* mRNA followed by five boxB loops, then the actin 3’-UTR. Three independent cell lines were selected for each CFB2 plasmid and CAT activity was measured with or without a 24h incubation with 100 ng/ml tetracycline to induce fusion protein expression (Supplementary Fig 9).

### The action of CFB2 depends on a conserved 16mer in the 3’-UTR of the *VSG* mRNA

*VSG* mRNA 3’-UTRs are relatively short, containing a CU-rich domain followed by 8-mer and 16-mer sequences that are conserved in almost all available *VSG* cDNA sequences (Supplementary Fig 10). Experiments using reporter mRNAs have shown that the 16-mer is required for high mRNA abundance and stability in bloodstream forms (12,13). It is also required for m^6^A modification of the poly(A) tail, which plays a role in *VSG* mRNA stabilization (67). To find out which sequences are responsive to CFB2, we integrated *GFP* reporter mRNAs with various 3’-UTRs (Fig 8A) into the tubulin locus, which results in constitutive RNA polymerase II transcription. As expected (13), the presence of the 16-mer resulted in relatively high expression in bloodstream forms (*GFP-VSG*), and its mutation to a scrambled version (*GFP-VSG m16*) decreased the GFP protein and mRNA levels to approximately the same level as the actin (GFP-ACT) control, while scrambling of the 8-mer *(GFP-VSGm8)* had no effect (Fig 8B, Supplementary Fig 11, Supplementary Fig 12).

**Figure 8.**
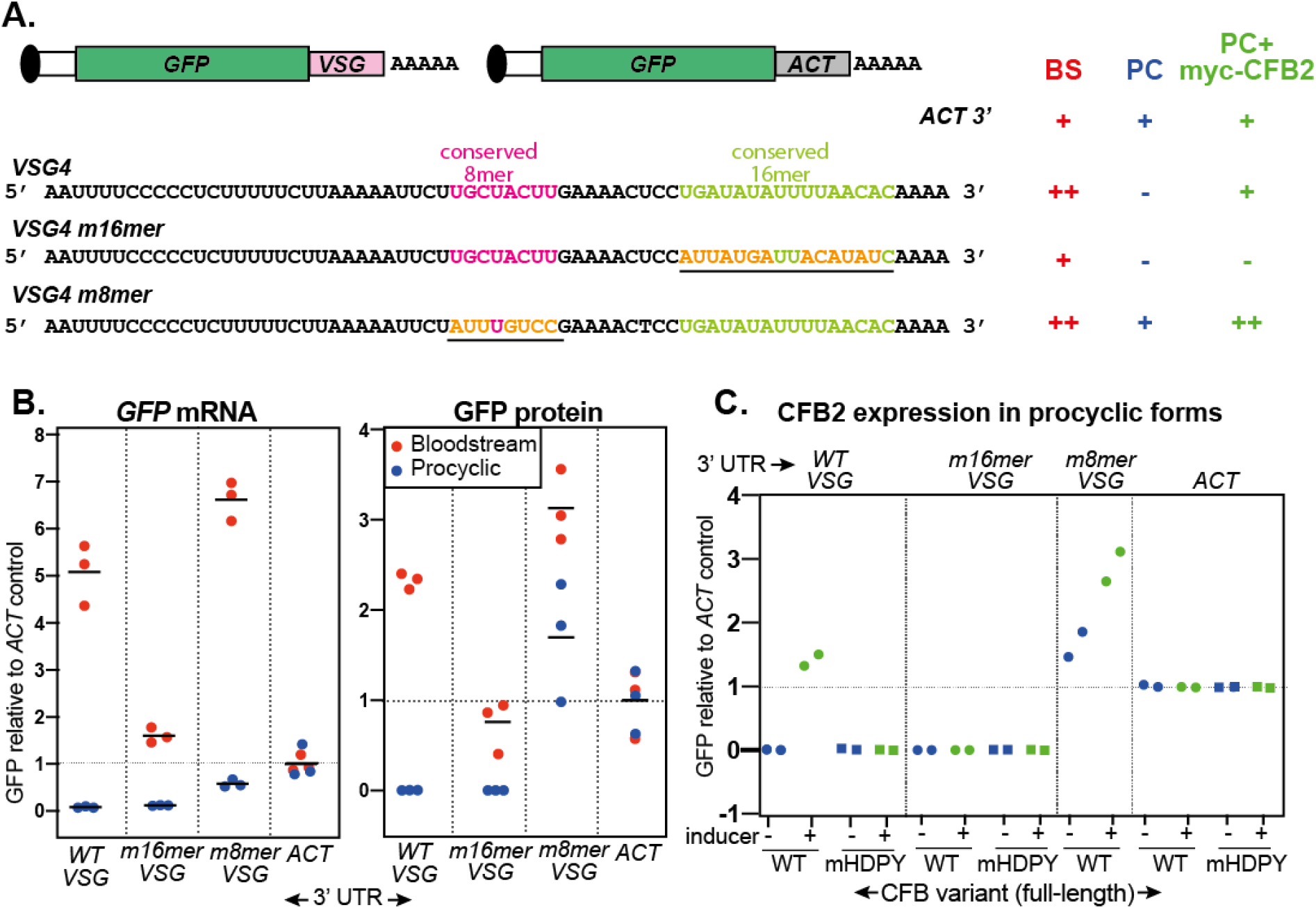
The role of the *VSG* 3’-UTR. **A.** Diagram of alternative reporters, and sequences of the 3’-UTRs. The 3’-UTR from the *VSG4* gene was cloned downstream of *GFP*, with polyadenylation directed by the *trans* splicing signal of the puromycin resistance cassette. After transfection of the linearized plasmid into trypanosomes, it integrates into the tubulin locus and is transcribed by RNA polymerase II. **B.** EATRO1125 *T. brucei*were transfected with the reporters as bloodstream forms, then three independent cloned lines for each construct were transformed to procyclic forms. Levels of GFP were quantified by Western blotting (Supplementary Fig 11), and of mRNA by Northern blotting (Supplementary Fig 12). All mRNAs were of the expected size. Results were normalized to the average for three controls expressing *GFP* mRNA with an actin *(ACT)* 3’-UTR.The left-hand panel shows *GFP* mRNA expression, and the right-hand panel, GFP protein. Individual measurements are shown, the bar is the arithmetic mean. **C.** Effect of CFB2-2xmyc expression on GFP expression in procyclic forms. For each GFP reporter from (B), we selected the procyclic clone with median expression. This was transfected with CFB2 inducible expression constructs 2 and 3 from Fig 7A and two independent cell lines were selected for each CFB2 plasmid. GFP protein was measured after 24h incubation with 100 ng/ml tetracycline, and quantified by Western Blot (Supplementary Fig 13) and results were normalized to the average for four controls expressing *GFP* mRNA with an actin (*ACT*) 3’-UTR.

In procyclic forms, CFB2 is not expressed. Upon transformation of the cloned trypanosomes to the procyclic form, the only *VSG* reporter giving detectable expression was that in which the 8-mer had been mutated (GFP-m8) (Figure 8B, Supplementary Fig 11, Supplementary Fig 12). This result implicates the 8mer in ensuring the low abundance of any *VSG* mRNAs that are made in the procyclic form.

We next inducibly expressed C-terminally myc-tagged CFB2 (CFB2-myc) in procyclic forms expressing the different reporters. As a control we expressed CFB2-myc with a mutated MKT1-interaction domain. These proteins were easier to detect than in bloodstream forms, suggesting that the auto-regulation has a degree of stage specificity (Supplementary Fig 13). Expression of CFB2-myc had no effect on GFP expression if the actin 3’-UTR was used (GFP-ACT), but caused marked increases in expression with the VSG 3’-UTR was used (GFP-VSG) (Fig 8C, Supplementary Fig 13). Expression of the reporter with the 8mer mutation (GFP-VSGm8) was also enhanced (Fig 8C, Supplementary Fig 13). In contrast, the reporter with mutant 16mer was unaffected by CFB2-myc expression (GFP-VSG m16) (Fig 8C, Supplementary Fig 13). As expected, the HDPY mutant of CFB2, which cannot interact with MKT1, had no effect on reporter expression. In some clones it was also equally expressed in the presence and absence of tetracycline, indicating a lack of selection for strong regulation, whereas the wild-type protein always showed cleanly inducible expression (Supplementary Fig 13).

These results show that CFB2 is able to increase expression from an mRNA bearing the VSG 3’-UTR and that this depends on the conserved 16mer sequence.

## Discussion

In this paper we describe a robust procedure to purify mRNAs and their associated proteins. Success of such procedures depends on their use to target mRNAs that have relatively high abundance; since we could identify proteins that were specifically associated with the alpha-tubulin mRNA, the threshold probably lies somewhere below 1% of total mRNA. The major limitation is the fact that mRNAs are about a thousand times less abundant than protein in cell lysates, so that a one-step purification is bound to be dominated by abundant protein contaminants. Our procedure could be used to purify sequentially several different abundant mRNPs from the same extracts; this is useful not only in saving material, but also because, if the order is varied, the different purifications act as internal specificity controls.

The role of the 3’-UTR in *VSG* mRNA abundance was demonstrated a quarter of a century ago (12), and conservation of 3’-UTR sequences was reported even earlier (68). It was also known that the action of the 3’-UTR is stage specific (12), and that *VSG* mRNA is degraded during differentiation to the procyclic form (69). By purifying the *VSG* mRNA together with its associated proteins, we have now identified CFB2 as the protein responsible for *VSG* mRNA retention in bloodstream forms. Moreover, we could demonstrate the mechanism by which CFB2 acts: recruitment of a stabilizing complex that includes MKT1, PBP1, PABP2 and the cap-binding translation initiation complex EIF4E6/G5. CFB2 action depends on the presence of a conserved 16mer in the *VSG* mRNA 3’-UTR. This sequence is also required for m6A modification of the *VSG* poly(A) tail (67). In principle CFB2 might recognize either the 16mer or m6A. However, m6A is present on many mRNAs in addition to *VSG* (67), whereas CFB2 depletion resulted in highly specific *VSG* loss. We therefore suggest that CFB2 recognizes the 16mer.

Although *VSG* transcription is shut off in procyclic forms, the parasites appear to have a failsafe mechanism that ensures that any accidentally produced *VSG* mRNA is rapidly degraded. We find here that a conserved 8mer sequence in the *VSG* 3’-UTR is implicated in this. Loss of *VSG* mRNA stability during differentiation is therefore the result of a combination of two processes: the disappearance of CFB2, and the expression of another, as yet unknown, protein that binds to the 8mer, represses translation and causes mRNA destruction.

Humans have at least 15 proteins that contain both RNA-binding and E3 ligase domains (70); they are implicated in autoubiquitination, and in either mono-ubiquitination or polyubiquitination of other targets (71–76). Our results suggest that the interaction of CFB2 with SKP1 promotes CFB2 degradation, presumably through auto-ubiquitination. This may fine-tune CFB2 abundance to limit VSG synthesis to an ideal level, preventing secretory pathway overload. SKP1 was detectable in purified MKT1-containing complexes (48,49), so CFB2 may be able to interact with SKP1 and MKT1 simultaneously. We do not know whether CFB2 has additional ubiquitination targets. These interactions, as well as the mode of CFB2 mRNA binding, require further investigation.

Adequate levels of *VSG* mRNA can be obtained only if the gene is transcribed by RNA polymerase I. In this paper we have shown, that in addition, the interaction between *VSG* mRNA and an mRNA binding-protein, CFB2, plays a crucial role in maintaining *VSG* mRNA abundance.

## Acknowledgements

We thank J. Gerst (Weizmann Institute) for providing the pcDNA4-MS2-CP-GFP-SBP plasmid and A. Gkeka and F. Aresta-Branco (DKFZ) for providing the cell line expressing the 3’-UTR of VSG2 and assistance in the generation of the tagged VSG construct. Medium was prepared by Ute Leibfried and Claudia Helbig (ZMBH). We thank Gloria Rudenko (Imperial College, London, UK) for the selectable VSG2 cell line and Kevin Leiss (ZMBH) for assistance in formatting additional genomes for use with Tryprnaseq. VSG 3’-UTR mutant cell lines were made and characterized by Meike Torwort and Lara Ruland during their BSc thesis work, and some CFB2 expression plasmids were made by Rachel Williams. We are indebted to Luise Krauth-Siegel (BZH) for allowing us to share her laboratory, including all equipment, after a disastrous flood in the ZMBH, and to Natalie Dirdjaja for helping us to move in. Electron microscopy was done by Charlotta Funaya of the Electron Microscopy Core Facility, Heidelberg University; Mass spectrometry was done in ZMBH facility under the leadership of Thomas Ruppert, and RNA Sequencing was done by David Ibbersson of Bioquant, University of Heidelberg. We thank Frauke Melchior (ZMBH) and especially Georg Stoecklin (University of Mannheim and ZMBH) for useful discussions and suggestions. Grant applications by CC to support this project were rejected by the ERC and the DFG.

## Methods

### *T. brucei* growth and manipulation

Bloodstream-form Lister 427 or EATRO1125 *T. brucei* expressing the *tet* repressor (pHD1313 or 2T1 cells) (77,78), were grown in HMI-11 medium (PAN Biotech). Cells with selectable VSG2 expression were a kind gift from Gloria Rudenko (Imperial College). These are ‘single marker’ derivative cell line carrying a puromycin resistance gene incorporated immediately behind the promoter of the active VSG2 ES (SM221) (79). Puromycin, phleomycin, hygromycin, G418 and blasticidin were used at 0.2, 0.2, 5, 2 and 5μg.ml^-1^ respectively for selection of recombinant clones. Regulation by the *VSG* 3’-UTR was investigated using EATRO1125 bloodstream-form trypanosomes, which were converted to procyclic forms by incubation with cis-aconitate and transfer to 27°C as previously described (43).

### Plasmids

Some cloning reactions were carried out with NEBuilder HiFi DNA assembly cloning kit (NEB) while others were made by conventional means. The plasmids are listed in Supplementary. Table S1. For RNAi of *CFB2*, a specific attB-tagged gene fragment (Tb927.1.4650, 378 bp) was amplified and cloned into pGL2084 (80) by Gateway recombination. The streptavidin-binding protein (SBP) tagged lambda-GFP construct was generated by amplifying the lambda-GFP and SBP coding regions in pHD2294 and pcDNA4-MS2-CP-GFP-SBP (a gift from J. Gerst - Weizmann Institute) respectively into *Hind* III and *Bam* HI sites of the pRpa vector (81). For overexpression of CFB2, we cloned the full ORF into the pRpa plasmid to add 6 C-terminal myc tags. We then subcloned the CFB2-6xmyc ORF into pHD1991, which drives constitutive expression and is inserted into the RRNA locus (82). For aptamer-tagging at the VSG2 native locus, a 225 bp fragment containing 5 *boxB* repeats in pHD2277 (66) was amplified and cloned into a derivative pSY37F1D-CTR-BSD plasmid (83) containing the *VSG2* CDS and 3’-UTR sequence to generate a *VSG2* gene with a boxB immediately after its endogenous 3’-UTR. Before transfection, the plasmid was linearized with *Bgl* II. Regulation by the *VSG* 3’-UTR was investigated using plasmids containing the *GFP* coding region and a puromycin selection cassette.

Prior to transfection into trypanosomes, the RNAi and CFB2 C-terminal myc-tag overexpression cassettes were digested with *Asc* I; all plasmids with pHD numbers were linearized with *Not* I. Linearized constructs were transfected into 2T1 cells (78) or cells containing integrated pHD1313 (77).

### RNA antisense purification-mass spectrometry (RAP-MS)

To identify proteins specifically interacting with *VSG2* and alpha tubulin (*TUBA*) mRNAs, we UV-cross-linked RNA and proteins *in vivo*, and captured RNA using biotinylated oligonucleotides using a protocol modified from (84). To reduce the costs associated with growth media we captured the two target RNAs in successive cycles from the same sample, exchanging the order of the probes to equalize the chance for contaminants. Mass spectrometry was done on pooled samples from 4 biological replicates (two each for each order), and the entire procedure was performed three times.

#### UV cross-linking

Cells at mid-log phase were collected (1,100g for 5 min), re-suspended into vPBS (PBS supplemented with 10 mM glucose and 46 mM sucrose, pH 7.6), then UV cross-linked on ice using two rounds of 0.18 J/cm^2^ of UV at 254 nm (Analytik Jena). Cells were collected, washed once with PBS, and pellets were flash-frozen in liquid nitrogen for storage at −80 °C.

#### Total cell lysate preparation

We lysed batches of ~9×10^9^ cells by completely re-suspending frozen cell pellets in 25 ml ice cold urea-based cell Hybridization Buffer (25 mM Tris pH 7.5, 500 mM LiCl, 0.25% dodecyl maltoside, 0.2% sodium dodecyl sulphate, 0.1% sodium deoxycholate, EDTA 5 mM, TCEP 2.5 mM and 4M urea). Next, the cell sample was passed ~10 times through a 27-gauge needle attached to a 25 ml syringe in order to disrupt the pellet and shear genomic DNA. At this point lysates were kept at −80 °C or incubated at 65 °C for 10 min before clearing by centrifugation for 10 min at 10,000g.

For the first purification, frozen cell pellets were resuspended in 8 ml ice cold detergent-based cell Lysis Buffer (10 mM Tris pH 7.5, 500 mM LiCl, 0.5% dodecyl maltoside, 0.2% sodium dodecyl sulphate, 0.1% sodium deoxycholate, 1 × Protease Inhibitor Cocktail EDTA-free, and 900 U of Murine RNase Inhibitor (New England Biolabs), and cell sample passed 10-15 times through a 27-gauge needle attached to a 25 ml syringe in order to disrupt the pellet and shear genomic DNA. The samples were then treated for 10 min at 37 °C adding 1X DNAse salt solution (2.5 mM MgCl2, 0.5 mM CaCl2), and 900 U of DNase I (Roche) to digest DNA. Samples were returned to ice and the reaction was immediately terminated by the addition of 16 ml 1.5X cold Hybridization Buffer to stop reaction.

#### RNA antisense purification of crosslinked complexes

1 ml of hydrophilic streptavidin magnetic beads (New England Biolabs) were washed 5 times with equal volume of Hybridization Buffer. Lysate samples were pre-cleared by incubation with the washed magnetic beads at 37 °C for 30 min with intermittent shaking at 1,100 r.p.m. on an Eppendorf Thermomixer C (30 s mixing, 30 s off). Streptavidin beads were then magnetically separated from lysate samples using a Dynal magnet (Invitrogen). The beads used for pre-clearing lysate were discarded and the lysate sample was transferred to fresh tubes twice to remove all traces of magnetic beads.

For each mRNA, we used seventeen 90-mer 5’-biotinylated complementary DNA oligonucleotides that spanned the entire length of the target RNAs. The oligonucleotides were heat-denatured at 85 °C for 3 min and then snap-cooled on ice. Probes and pre-cleared lysate were mixed and incubated at 65°C using an Eppendorf thermomixer with intermittent shaking (30 s shaking, 30 s off) for 1.5 h to hybridize probes to the target RNA. Samples were then incubated with washed C1 Streptavidin coated magnetic beads (Thermo Fisher Scientific) at 65 °C for 2.5h on an Eppendorf Thermomixer C with intermittent shaking as above. Beads with captured hybrids were washed 5 times with Hybridization Buffer at 65 °C for 5 min to remove non-specifically associated proteins. Then, samples were washed twice with DNAse Buffer (50 mM Tris pH 7.5, 300 mM LiCl, 0.5% NP40, 0.1% NLS and 1X DNAse salt solution) and incubated with intermittent shaking (30 s shaking, 30 s off) for 15 min with 15 units of DNAse I (Roche) to remove DNA traces. Around 2% of the total beads were removed and transferred to a fresh tube after the final wash to test RNA capture by RT-qPCR. The remaining beads were resuspended in Benzonase Elution Buffer (20 mM Tris pH 8.0, 2 mM MgCl2, 0.05% NLS, 0.5 mM TCEP) for subsequent processing of the protein samples.

#### Elution of protein from the beads

Elution of captured proteins from streptavidin beads was achieved by digesting all nucleic acids with 125 U of Benzonase nonspecific RNA/DNA nuclease for 2 h at 37 °C. Beads were then magnetically separated from the sample using a Dynal magnet (Invitrogen); the supernatant containing eluted proteins was first partially concentrated in a Speedvac to about 200 μl and finally methanol/chloroform precipitated (85).

#### Elution and analysis of RNA samples

Beads with hybrids were magnetically separated using a Dynal magnet and the supernatant was discarded. Beads were then resuspended by pipetting in 50 μl NLS RNA Elution Buffer (20 mM Tris pH 8.0, 10 mM EDTA, 2% NLS, 2.5 mM TCEP). To release the target RNA, beads were heated for 2 min at 95 °C. Beads were then magnetically separated and the supernatants containing eluted target RNA were digested by the addition of 1 mg ml^-1^ Proteinase K for 1 h at 55 °C to remove all proteins. The remaining nucleic acids were purified using the RNA Clean & Concentrator Kit (Zymo).

For synthesis of cDNA, the Maxima First Strand cDNA Synthesis Kit for RT-qPCR (Thermo Fisher) was used following manufacturer’s protocol. To quantify RNA enrichment, real time PCR was performed in triplicate using Luna^®^ Universal qPCR Master Mix (New England Biolabs) with variable amounts of cDNA and 0.5 μM of target-specific primers in a CFX connect instrument (Bio-Rad). Primer sequences are shown in Supplementary Table S1.

#### Mass spectrometry analysis

Proteins from four independent purifications (material from about of 3×10^10^ trypanosomes, two with tubulin mRNA selected first, and two with *VSG*) were pooled, subjected to denaturing SDS-PAGE, and analyzed by the ZMBH Core facility for mass spectrometry and proteomics as previously described (26). Data were quantitatively analyzed using Perseus (86).

### RNA-binding protein purification and identification (RaPID)

Rapid experiments and controls. To identify CFB2 specifically interacting with VSG2 mRNA, we performed captures of lambda-GFP-SBP proteins in cells expressing boxB-tagged *VSG2* mRNA and lambdaN-GFP-SBP. Cells lacking either component were used as controls.

#### Formaldehyde crosslinking and cell lysis

Cells at mid-log phase were collected, re-suspended in PBS and then cross-linked with 0.01% formaldehyde at room temperature for 10 min with slow shaking. The cross-linking reaction was terminated by adding 1 M glycine-NaOH buffer (pH 8.0) to a final concentration of 0.125 M and additional shaking for 2 min. The cells were then washed once with ice-cold PBS buffer and the pellet was flash-frozen in liquid nitrogen, and stored at −80°C. Cell pellets were thawed by addition of ice-cold lysis Buffer (25 mM Tris-HCl pH 7.5, 175 mM KCl, 0.5% NP40, 1 mM DTT, and 120 U RNAse inhibitor (New England Biolabs) supplemented with EDTA-free protease inhibitors (Roche). Samples were then subjected to three cycles of sonication (10 pulses of 0.5 s) followed by 1 min rest between cycles at 4 °C (Branson Ultrasonics Sonifier S-250). The extract was then supplemented with 1x DNAse salt solution and 100 U DNAse I (Roche) and incubated for 30 min at 4 °C on rotator. Samples were returned to ice and the reaction was immediately terminated by the addition of 10 mM EDTA and 10 mM EGTA. The extract was finally clarified by centrifugation for 20 min at 15,000 × g, the supernatant removed to a new microcentrifuge tube and protein concentration determined using the Bradford assay (Bio-Rad).

#### Precipitation of RNP complexes

RaPID was performed essentially as described by (87) with a few modifications. In order to block endogenous biotinylated moieties, the protein aliquot taken for pull-down was incubated with 10 mg of free avidin (Sigma) per 1 mg of protein input at 4°C for 1 h with constant rotation. In parallel, streptavidin magnetic beads (New England Biolabs) were washed 5 times with 1 ml of Lysis Buffer supplemented with EDTA 5mM. Pull-down was then performed by adding the indicated amount (see figure legends) of avidin-blocked total cell lysate to the beads, followed by incubation at 4°C for 3 h with constant rotation. Beads with captured mRNPs were washed 3 times with lysis Buffer and three times with washing buffer (25 mM Tris-HCl pH 7.5, 300 mM KCl, 0.5% NP40, 1 mM DTT, 5 mM EDTA and 40 U/ml RNAse inhibitor (New England Biolabs) at 4 °C for 5 min to remove non-specifically associated proteins. For elution of the cross-linked RNP complexes, 250 μl of washing buffer supplemented with 6 mM free biotin (Sigma) was added to the beads, followed by 1 h of incubation at 4°C with rotation. To reverse the crosslink for RNA analysis, samples were incubated at 70°C for 1 h with cross-link reversal buffer (50 mM Tris-HCl pH 7.5, 5 mM EDTA, 10 mM DTT, and 1% SDS).

### Protein blotting and antibodies

Protein samples were run according to standard protein separation procedures, using SDS-PAGE. The following primary antibodies were used: mouse α-GFP (1:2000), mouse α-myc (1:2000), rat α-RBP10 (1/500), rabbit α-aldolase (1/2000). We used horseradish peroxidase coupled secondary antibodies (α-mouse and α-rabbit, Bio-Rad, 1:2000). Blots were developed using an enhanced chemiluminescence kit (Amersham) according to the manufacturer’s instructions. Densitometry was performed using Fiji v. 2.0.0.

### Immunoprecipitation of mRNA-protein complexes

For immunoprecipitation, cells expressing CFB2-6xmyc were first crosslinked using either UV irradiation or formaldehyde as described above. In some cases, MG132 was added to a final concentration of 10 μg/ml for 1 h before crosslinking.

Formaldehyde-treated cell extracts were made as described for RaPiD.

UV irradiated cells were washed in cold PBS and the cell pellet snap frozen in liquid nitrogen or used immediately. Cells were lysed by resuspension in lysis buffer (50 mM Tris-HCl, pH 7.5, 2 mM MgCl2, 10 mM KCl, 0.1 mM DTT, 0.5% (w/v) NP-40, 100 U/ml murine RNase inhibitor (NEB), and EDTA-free protease inhibitor cocktail (Roche)). Next, samples were passed ~10 times through a 27-gauge needle in order to disrupt the pellet and shear genomic DNA. After adding KCl to 150 mM, the lysate was centrifuged at 4°C for 10 min at 12,000 x g.

The supernatants from both types of extract were subjected to immunoprecipitation using anti-c-Myc magnetic beads (Pierce) for 3 h at 4°C. After extensive washing in lysis buffer supplemented with the indicated amount of KCl (see figure legends), RNA was extracted from the immunoprecipitated material using Trifast reagent (Peqlab, GMBH).

### RNAi by direct dsRNA transfection

The template for *CFB2* dsRNA production was identical to the one employed in the tet-inducible system, and was amplified using primers that include a T7 promoter for *in vitro* transcription (Supplementary Table S1). dsRNA was made using the MEGAscript RNAi Kit (Ambion, Thermo Fisher Scientific). 20 μg of dsRNA or the same buffer volume (for the mock control) were transfected into 30 million cells, using the X-001 program of an AMAXA Nucleofector.

### Electron Microscopy

For transmission electron microscopy, samples were prepared exactly as described in (88) after 16 h of CFB2 RNAi induction. The blocks were sectioned using a Leica UC6 ultramicrotome (Leica Microsystems Vienna) in 70 nm thin sections. The sections were placed on formvar coated grids, post-stained and imaged on a JEOL JEM-1400 electron microscope (JEOL, Tokyo) operating at 80 kV and equipped with a 4K TemCam F416 (Tietz Video and Image Processing Systems GmBH, Gautig).

### RNASeq

RNA was prepared after 9h of *CFB2* RNAi induction in cells expressing *VSG2*. rRNA was depleted using complementary oligonucleotides and RNaseH as previously described (89). It was fragmented, and cDNA libraries were prepared using an Illumina kit before sequencing (MiSeq). Raw reads trimmed then aligned to *T. brucei* genomes using TrypRNASeq (90) and genomes downloaded from TritrypDB. To assess levels of transcripts from expression sites, reads were aligned to the 2018 version of the Lister 427 genome (3). Each read was allowed to align once (such that reads for repeated genes are distributed among the copies) and these results are in Supplementary Table S3. To obtain more information about annotation of any affected chromosome-internal genes, we repeated the alignment using the well-annotated TREU927 genome (91) (Supplementary Table S4). To look for changes in expression, we used DeSeqU1 (92), a user-friendly RStudio DeSeq2 (93) application. To look for enrichment of particular functional classes, we considered a list of genes in which each individual sequence is present only once (that is, additional gene copies have been removed) (94). Since, in this list, reads from repeated genes will be under-represented, we multiplied each read count by the gene copy number. Copy numbers were obtained by repeating the alignments, allowing each read to align twenty times.

The distribution of reads across Lister427 chromosomes was visualized using Artemis. For this, we chose induced and uninduced samples with similar total aligned read counts.

### CFB2 expression in *E. coli*

For expression of CFB2 in *E. coli*, the coding sequence of CFB2 was cloned into pET-NusA for N-terminal tagging with a 6 × His-tag and the transcription termination/antitermination protein NusA, which was then transformed into BL21 competent *E.coli*. Individual transformants were inoculated in 5 mL of LB^Kan^ each, and grown for 16 h at 27 °C. After that, OD_600_ measurements were performed every hour, until the cultures reached an OD_600_ of 0.8. For each inoculum, 1 mL was collected by centrifugation (2 min, 16,000 × g), and the supernatant was subsequently removed. The pellet was then lysed in Laemmli buffer (“uninduced” sample) and stored at −20 °C until further use. Recombinant protein expression was induced by addition of IPTG at a final concentration of 1 mM to the remaining 4 mL of the culture. Furthermore, another 1 mL of LB^Kan^ supplemented with IPTG (1 mM) was added, after which the samples were incubated at 27 °C for 14 h. Upon reaching an OD_600_ of 2.5-3, 300 μL were collected per sample by centrifugation at 16,000 × g for 2 min. These “induced” samples were processed and lysed similar to the “uninduced” samples. For analyzing protein expression, the samples were boiled at 95 °C for 10 min and subsequently separated using 10% SDS-polyacrylamide gels, which were then stained by Coomassie staining.

CFB2 was also expressed as a fusion with 6 his tags and the GB protein, by cloning into pETGB. Expression was induced and inclusion bodies purified as above. This protein was used for rabbit-antisera purification and as a positive control on Western blots.

### Analysis of soluble and insoluble *E. coli* fractions

Induction and harvest of the cultures were performed as described above. However, instead of lysing the cells with Laemmli buffer, the pellet was resuspended in 350 μL of bacterial lysis buffer (50 mM Tris [pH 7.5], 100 mM NaCl, 1 mM EDTA, 1 mM DTT, 1 μg/mL aprotinin, and 1 μg/mL leupeptin), after which 25 μL of lysozyme solution (stock: 10 mg/mL in Tris-HCl [pH 8.0]) were added. After mixing the samples by vortexing for 3 sec, lysis was performed for 4 h at 4 °C with constant rotation. This was followed by 3 freeze/thaw cycles using liquid nitrogen and centrifugation for 10 min at 16,000 × g and 4 °C. Soluble (supernatant) and insoluble (pellet) fractions were analyzed separately by SDS-PAGE and Coomassie staining of the gels.

### Purification of inclusion bodies for antibody production

For large scale purification of inclusion bodies, 1 L of an IPTG-induced culture was grown at 27 °C until reaching an OD_600_ 0.8, and harvested by centrifugation at 5000 rpm for 20 min. The pellet was washed once with 1 × PBS and stored at −80 °C until further processing. Lysis of the bacteria was performed by addition of 500 μL of bacterial lysis buffer and 50 μL of lysozyme solution (see above). Furthermore, the samples were mixed by vortexing for 3 sec, incubated for 4 h at 4 °C with constant rotation, and subsequently subjected to 3 freeze/thaw cycles using liquid nitrogen. Inclusion bodies were pelleted by 20 min centrifugation at 15,000 rpm and 4 °C, washed once with bacterial lysis buffer containing 1% TritonX-100, and eventually resuspended in 10 mL of 8 M urea. These were then left to dissolve for 16 h at 4 °C with constant rotation.

After determining the protein concentration spectrophotometrically (A280; Nanodrop™, Thermo Fisher Scientific™, Karlsruhe, Germany), 0.5 mg of the protein in inclusion bodies was then separated by SDS-PAGE and the band corresponding to NusA-CFB2 was cut from the gel. The gel slice was submitted to David’s Biotechnology (Regensburg, Germany) for the generation of antisera in rabbit.

### Purification of anti-CFB2 antibodies

Polyclonal antisera were purified by adsorption of total rabbit serum to immobilized recombinant CFB2-Gb1 Carrier protein. 100 μg of the CFB2 recombinant protein was resolved in SDS-PAGE gel and transferred to PVDF membrane. CFB2 containing region was excised, fragmented in small pieces and transferred to a 1,5 ml tube. The membrane was blocked with 5% milk in PBS/0,05% tween for 1 hour at RT with agitation, washed twice in PBS containing 0,05% Tween-20 and incubated with 1 ml of antisera for 16 hours at 4°C with agitation. After incubation, the depleted antiserum was discarded and the membrane washed 3 times in PBS/0,05% Tween-20. Finally, the bound antibodies were eluted with 200 μl of 0.1 M glycine pH 2.4 for 5 min, under vortex and neutralized with 20 μl of Tris-HCl pH 8.0.

### Tethering assays

Tethering assays were done using cells expressing mRNAs with a chloramphenicol acetyl-transferase open reading frame and a truncated version of the trypanosome *actinA* 3’-UTR, with 5 boxB sequences immediately upstream of the 3’-UTR (95). Lines with tetracycline-inducible expression of different versions of CFB2 (as shown in Figure 7) were made, and CAT was assayed with or without tetracycline addition.

## Notes

### Competing Interest Statement

The authors have declared no competing interest.

