## Supplemental Figures & Legends for "The trypanosome Variant Surface Glycoprotein mRNA is stabilized by an essential unconventional RNA-binding protein"

### Supplementary Figures

#### Supplementary Fig 1

**CFB2 domain conservation and evolution**

A. Evolutionary trees, made with CLUSTALW, for different portions of CFB proteins from different Kinetoplastids. Tb927.1.4650 was chosen as representative of CFB1 and Tb927.1.4540 is CFB2. *T. vivax* genes shown are TvY486_0102030 (Tv30), TvY486_0102040 (Tv40), TvY486_0404500 (Tv45), Tv486_0102050 (Tv50) and TvY486_0011670 (Tv67). *T. congolense* genes are TcIL3000_1_1860 (Tco60) and TcIL3000_1_1870 (Tco70). Others are TRSC58_03601 (*T. rangeli*, Tra) TM35_000322060 (*T. theileri*, Tth); TcCLB.506529.80 (*Trypanosoma cruzi*, Tcr); PCON_0072200 (*Paratrypanosoma confusum*, Pcon), DQ04_06951020 (*Trypanosoma grayi*, Tgr).

#### Supplementary Fig 2

**Effects of CFB2 RNAi and uncropped images for Figures 2 and 4**

A. Uncropped images for Figure 2

B. Cells expressing *VSG2* and CFB2-6xmyc were used. Some were pre-incubated with MG132, with or without UV irradiation; others were mildly fixed with formaldehyde. After lysis and immunoprecipitation, the amounts of *VSG2* mRNA and rRNA were measured by reverse transcription and real-time PCR. The ratios were then compared to those in the input extracts. VSG2 mRNA was enriched in 5/6 experiments.

C. The cells used expressed *VSG3* with the *VSG2* 3'-UTR, from the active expression site. The Figure shows relative quantification of RNA transcript levels corresponding to *VSG3* and alpha-tubulin (*TUB*) mRNAs detected at different sampling time-points (3, 6, 9h) after induction of *CFB2* RNAi. CTR (co-transposed region) is a sequence located upstream of the *VSG* gene, which is transcribed from the same promoter but present only in the mRNA precursor. Results were normalised against 18S. This is a single experiment. The decreases in all mRNAs after 9h may have been caused by decreasing cell viability.

D. Northern blot analysis corresponding to panel (C).

E. Double-stranded CFB2 RNA was introduced into trypanosomes naturally expressing *VSG22* by electroporation. Cells that were electroporated with no dsRNA served as the control. RNA was prepared 3h and 6h after electroporation, and levels of *CFB2, VSG22* and alpha-tubulin mRNA were assayed by reverse transcription and real-time PCR. *CFB2* mRNA was reduced within 3h, and a marked reduction in *VSG22* mRNA had occurred after 6h. There was a reduction in *TUB* also, but it was not so marked.

F. CFB2 RNAi for 8h does not cause a general suppression of translation. Trypanosomes expressing VSG2 and with inducible RNAi targeting CFB2 were used. Tetracycline was added for the times shown. Cells were then harvested, washed, and incubated with [$^{35}$S]-methionine for 20 min. Afterwards the reactions were run on a polyacrylamide gel, which was dried before detection. Parallelly and before drying, one lane was blotted and probed with anti-VSG2 antibodies. The stained gel is above and the signal from [$^{35}$S]-methionine is shown below. The prominent band just below 50 KDa is probably tubulin (TUB), the ones above it are probably VSG22 and VSG2 naturally expressed in the 1313 and 2T1 cell lines, respectively. Note that VSG22 runs slightly slower than VSG2. After 7.5h the VSG2 band is much less prominent.

G. Uncropped images for Fig 4B.

**Supplementary Fig 3**

**Roles of the F-box and HDPY sequence in protein-protein interactions**

Yeast 2-hybrid assays were done using CFB2 with critical residues of the F-box or MKT1 interaction motif mutated to alanine, as indicated in Figure 3A.

A. Growth of yeast expressing the proteins as indicated; the upper panel group shows growth under selection for the interaction (minus tryptophan, leucine, alanine and histidine) and the lower panel shows selection only for the presence of the plasmids.

B. Expression of the fusion proteins. The table indicates which lane is which.

**Supplementary Fig 4**

**Effects of CFB2 RNAi on the transcriptome**

Reads were aligned to the Lister 427 (2018) genome.

A. Principal component analysis.

B. Correlations between replicates, and between the parental cell line and the RNAi cell line without tetracycline. Every spot represents a single open reading frame.

**Supplementary Fig 5**

**Effects of CFB2 RNAi on ultrastructure - additional images**

Cells were fixed and stained after 16h in tetracycline.

A. Cell with a large flagellar pocket and numerous cytosolic axonemes and paraflagellar rods. The enlargement shows that the VSG layers on the surface and flagellar pocket appear similar.

B. The upper cell has a normal flagellar pocket (enlarged, a) as well as a cytosolic flagellum. The enlarged vesicle or flagellar pocket "b" appears to lack an electron-dense coat. The structures marked ‘?’ look like multivesicular bodies or autophagosomes; one of them contains part of an axoneme.

**Supplementary Fig 6**

**Effects of CFB2 RNAi on ultrastructure - additional images**

Cells were fixed and stained after 16h in tetracycline.

A. Cell containing a cytosolic invagination that includes three axoneme-PFR structures.

B. Full flagellum enclosed in membrane with underlying corset microtubules. There is also a set of four microtubules with underlying ER, characteristic of the flagellar attachment zone (FAZ) (1).

C. Cross-section of a similar tube containing a flagellum. The dots to the left are a flagellar attachment zone (1).

D. Cell with extended internal membranes with underlying corset microtubules. This was seen more rarely than the other abnormalities shown.

**Supplementary Fig 7**

**Antibody tests and CFB2 protein expression**

CFB2 was expressed as a fusion with six His tags and NusA at the N-terminus, in BL12 *E. coli*. Purified inclusion bodies were solubilized in 8M urea and subjected to denaturing SDS-PAGE. The band containing 6x-His-NusA-CFB2 was used to raise antibodies in a rabbit. For affinity purification, membrane-bound 6xHis-GB-CFB2 was used.

A. Coomassie-stained gel showing bacterial lysates, with or without induction of expression of either 6xHis-GB-CFB2 or 6xHis-Trx-CPSF73 (negative control). Serial dilutions of *E. coli* extracts are shown. A key for the different arrows is to the right at the top. The 100-fold dilution of 6xHis-GB-CFB2 expressing cells (lane 3) gives roughly the same CFB2 band intensity as the markers (probably 1-5 µg each, data not available). The 100-fold dilution exceeds the Coomassie blue detection limit of approximately 100ng.

B. CFB2 has relatively low abundance in bloodstream-form trypanosomes. The upper two panels are different exposures of a Western blot, and the third panel is the Ponceau red stained membrane. In lane 1, lysate of $2x10^6$ bloodstream forms expressing CFB2-6xmyc from an rRNA promoter was boiled in Laemmli buffer before loading; in lane 2, $10^7$ bloodstream forms without tagged CFB2 expression were used and the extract was not boiled. Lane 3 is a $10^{-3}$ x dilution of *E. coli* containing the 6xHis-GB-CFB2 plasmid, but not induced; lanes 4 is the same, but induced. Lanes 5-7 are successive 10-fold dilutions of the same extract. Lanes 8-11 are equivalent *E.coli* extracts, but the cells express 6xHis-Trx-CPSF73; larger amounts of input protein were used because the recombinant protein was poorly expressed (see panel A). Lane 12 is an extract of *S. cerevisiae* expressing AD-CFB2-myc; insufficient is present for detection. The top panel shows that the antibody interacts with 6xHis-GB-CFB2, and not with 6xHis-Trx-CPSF73, although there is a faint cross-reaction with a constitutively expressed *E. coli* protein of around 60 kDa. The purified antibody, which was raised against CFB2-NusA, is therefore interacting predominantly with CFB2 (trypanosomes do not contain NusA). The CFB2 band in lane 2 has approximately the same intensity as the $10^5$-fold dilution of 6xHis-GB-CFB2 *E. coli* lysate. From panel A, this dilution probably contains 1-10 ng of CFB2, which corresponds to between 1000 and 10,000 protein molecules per cell.

C. Purified anti-CFB2 antibody recognises a band of the expected size that decreases after *CFB2* RNAi and increases after proteasome inhibition. In lanes 1-6, $1x10^7$ cells were lysed in sample buffer and loaded onto the gel. In lane 6 and also lane 7, which contains a lysate of *E. coli* expressing recombinant protein (CFB2-GB), the samples were boiled before loading. Lanes 1 and 6 are wild-type trypanosomes that had been incubated with MG132 for 1h. For lanes 2-5, cells with inducible RNAi were used, incubated with tetracycline inducer for the times indicated. The upper image was obtained after the membrane had been incubated with affinity-purified antibody (1/1000 dilution) for 1h. For the central panel, the membrane was cut to remove the region above 56 kDa that contains a cross-reacting band, then incubated overnight with the antibody. The lowest panel is the blotted membrane stained with Ponceau red. The key to the different arrows is above the membrane panels.

D. Cells used expressed various fusion proteins as shown in the top diagram. Separate Western blots from the same gel were used, with $2x10^6$ loaded for detection of the myc epitope and $2x10^7$ for CFB2. The affinity-purified anti-CFB2 antibody was incubated with the membrane overnight. Arrows are as in panel A. The cells expressing CFB2-6myc express VSG2 and the remaining cells express VSG22.

E. The effect of MG132 does not depend on an intact CFB2 N-terminus. Cells constitutively expressing CFB2 with 6 myc tags at the N- or C-terminus were used, with or without a one-hour incubation with MG132. The Western blot was incubated with anti-myc antibodies. The slower-migrating cross-reacting band also serves as a loading control.

**Supplementary Fig 8**

**Tethering screen results and the conserved region C-terminal to the F-box**

A. CFB2 fragments lacking the F-box activated expression in a tethering screen. The cell line used expressed a blasticidin deaminase mRNA with five copies of boxB. A library that contained random ~1kb genomic fragments was transfected, and cells were selected with increasing amounts of blasticidin. Afterwards, the genomic fragments were amplified and sequenced in order to locate the N-termini of expressed fusion proteins. The graph shows the fold enrichment of sequences grown in 10x the normal level of blasticidin, relative to no blasticidin (2). High representation of a sequence indicates that a protein starting at that residue was able to increase blasticidin deaminase expression. Only fusion proteins lacking the F-box were active in the assay. A separate assay of complete open reading frames yielded no enrichment of CFB2 (3).

B. Alignment of the conserved domain downstream of the F-box. The colour code for residues is red: non-polar; green: polar; blue: acidic; purple: basic.

**Supplementary Fig 9**

**Expression of CFB2 lambdaN fusion proteins in bloodstream forms**

A. Diagram showing the different fusion proteins and their names: this is the same as the relevant part of Figure 7.

B. Western blots for $2x10^6$ cells with or without 24h induction with tetracycline. The blots were incubated together with anti-myc antibody, and the signals were detected together in order to have identical exposures. This panel shows that the C-terminal half of CFB2 is much easier to detect than the full-length protein or the N-terminal half.

C. As B: Mutation of the SKP1 binding site results in the N-terminal half becoming detectable.

D. As B: Mutation of the SKP1 binding site results in the full-length protein becoming readily detectable, whereas mutation of the MKT1 binding site has little effect on expression of either full-length CFB2 or the C-terminal half.

**Supplementary Fig 10**

Alignment of *VSG* 3'-UTRs, downloaded from Genbank. Nearly all of the sequences include the conserved C-rich region, 8mer (pink) and 16mer (green). Many of the sequences are from cDNA, and in the few cases where the conserved regions are changed, there might be cloning or sequencing error - for example the apparently mutant cDNA sequence of *VSG22* does not match the genomic copy. On the top two lines, the *VSG2* 3'-UTR is compared with that of *VSG4*, which was used for GFP reporter experiments; the colour code is the same as that in Figure 8.

**Supplementary Fig 11**

Western blots used for Figure 8.

Groups of two blots that were hybridised and exposed simultaneously are shown as different experiments. For bloodstream forms, quantification was done for experiment 2, except for VSG m16 clone 3, which was lost in that experiment. Instead, VSG m16 clone 3 GFP was quantified relative using the blot of experiment 1. For procyclic forms, data from experiment 2 were used. Signals were normalized to the Ponceau red stain.

**Supplementary Fig 12**

Northern blots used for Figure 8.

Two blots were made using the same RNA samples; quantification was done using the blots on the right. The faint band at 1.5kb is a cross-reacting RNA. Signals were normalized to the methylene blue stain.

**Supplementary Fig 13**

**Expression of different versions of CFB2-myc in cells with GFP reporters.**

Western blots with detection of myc and GFP are above and corresponding loading control (anti-S9) and Ponceau-stained membrane images are below. The migration positions of CFB2-2xmyc, GFP and S9 are indicated. The sequences of the *VSG* 3'-UTRs are shown in Figure 8.

A. *GFP* reporter mRNA with a wild-type *VSG* 3'-UTR.

B. *GFP* reporter mRNA with 16mer mutant *VSG* 3'-UTR.

C. *GFP* reporter mRNA with 8mer mutant *VSG* 3'-UTR.

D. *GFP* reporter mRNA with *ACT* 3'-UTR.

**Supplementary Tables**

**Supplementary Table 1**

**Plasmids and oligonucleotides used in the experiments.**

**Supplementary Table 2**

**Mass spectrometry results.**

The Legend is on the first sheet.

**Supplementary Table 3**

**Effects of RNAi on the transcriptome: alignment to the Lister427_2018 genome.**

Reads were aligned to the Lister427_2018 genome. Each read was allowed to align once. The legend is on the first sheet of the table. Annotations are often approximate since it is sometimes not clear which TREU927 gene is a true homologue of the Lister427 versions, especially for gene families.

**Supplementary Table 4**

**Effects of RNAi on the transcriptome: alignment to the TREU927 genome**.

Reads were aligned to the TREU927 genome. Each read was allowed to align once. The legend is on the first sheet of the table. The sheet that includes the unique gene set, in which each set of repeated genes has only one representative, allows judgement of whether any particular functional class is enriched.

Fig.1

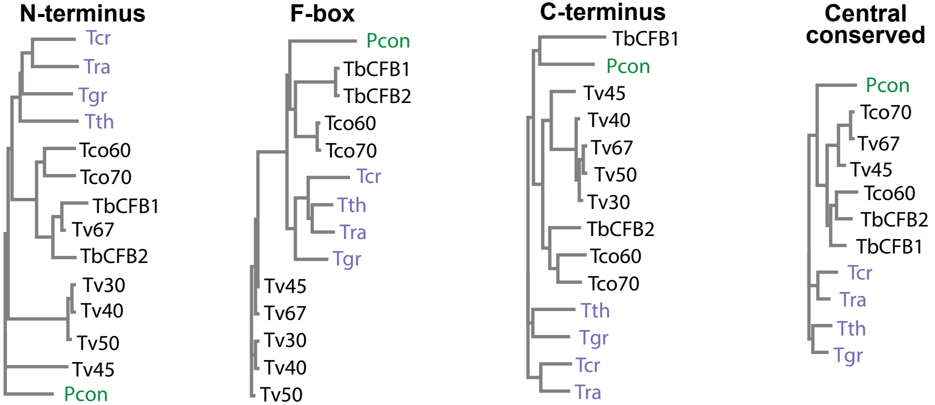
N-terminus
Tcr
Tra
Tgr
Tth
Tco60
Tco70
TbCFB1
Tv67
TbCFB2
Tv30
Tv40
Tv50
Tv45
Pcon
F-box
Pcon
TbCFB1
TbCFB2
Tco60
Tco70
Tcr
Tth
Tra
Tgr
Tv45
Tv67
Tv30
Tv40
Tv50
C-terminus
TbCFB1
Pcon
Tv45
Tv40
Tv67
Tv50
Tv30
TbCFB2
Tco60
Tco70
Tth
Tgr
Tcr
Tra
Central conserved
Pcon
Tco70
Tv67
Tv45
Tco60
TbCFB2
TbCFB1
Tcr
Tra
Tth
Tgr

Fig. 2

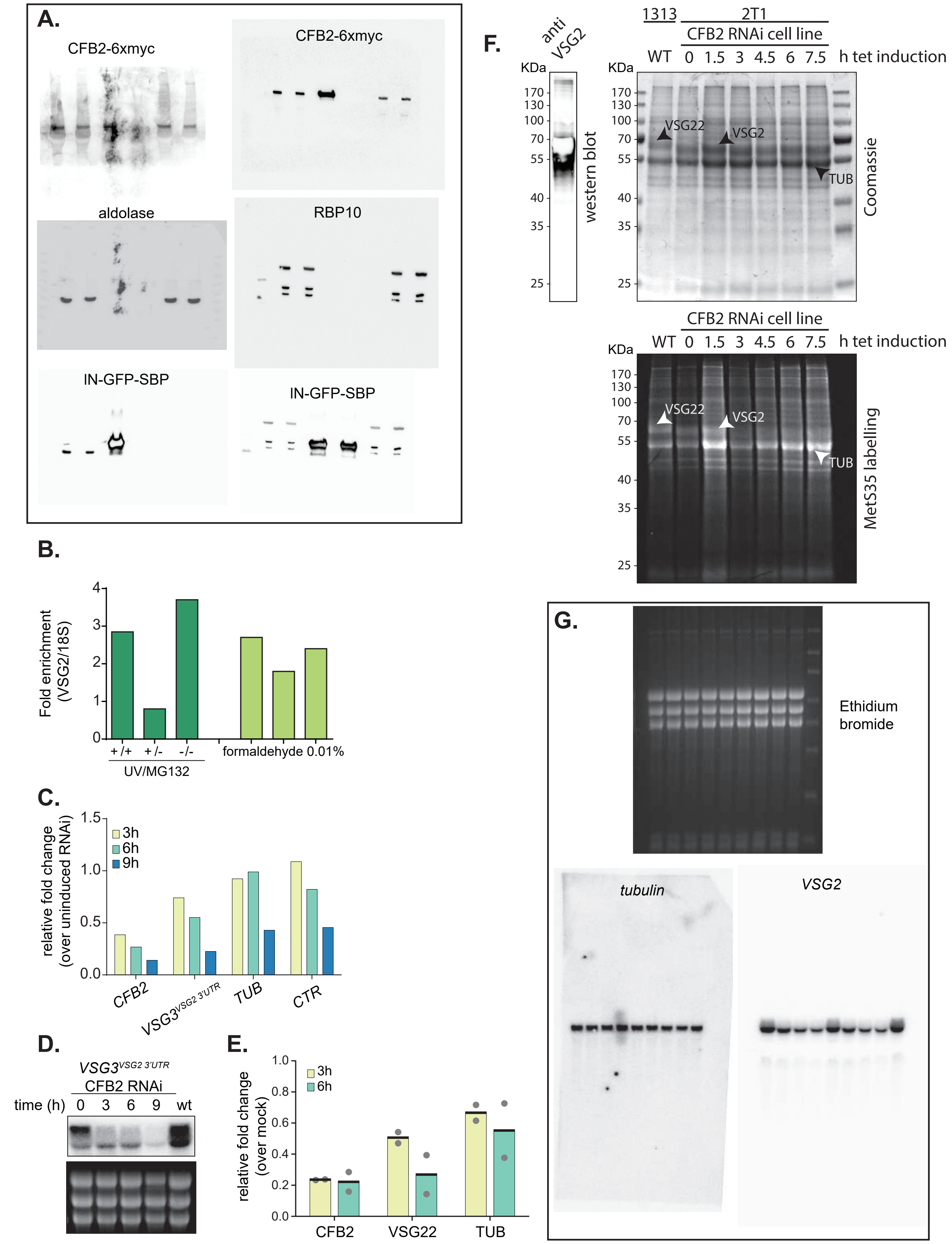
A.
CFB2-6xmyc
CFB2-6xmyc
aldolase
RBP10
IN-GFP-SBP
IN-GFP-SBP
B.
Fold enrichment (VSG2/18S)
+/+
+/-
-/-
UV/MG132
formaldehyde 0.01%
C.
relative fold change (over uninduced RNAi)
3h
6h
9h
CFB2
VSG3VSG2 3'UTR
TUB
CTR
D.
VSG3VSG2 3'UTR
CFB2 RNAi
time (h) 0 3 6 9 wt
E.
relative fold change (over mock)
3h
6h
CFB2
VSG22
TUB
F.
anti VSG2
KDa
170
130
100
70
55
40
35
25
western blot
1313
2T1
CFB2 RNAi cell line
WT 0 1.5 3 4.5 6 7.5 h tet induction
VSG22
VSG2
TUB
Coomassie
CFB2 RNAi cell line
WT 0 1.5 3 4.5 6 7.5 h tet induction
MetS35 labelling
G.
Ethidium bromide
tubulin
VSG2

Fig. 3

A.

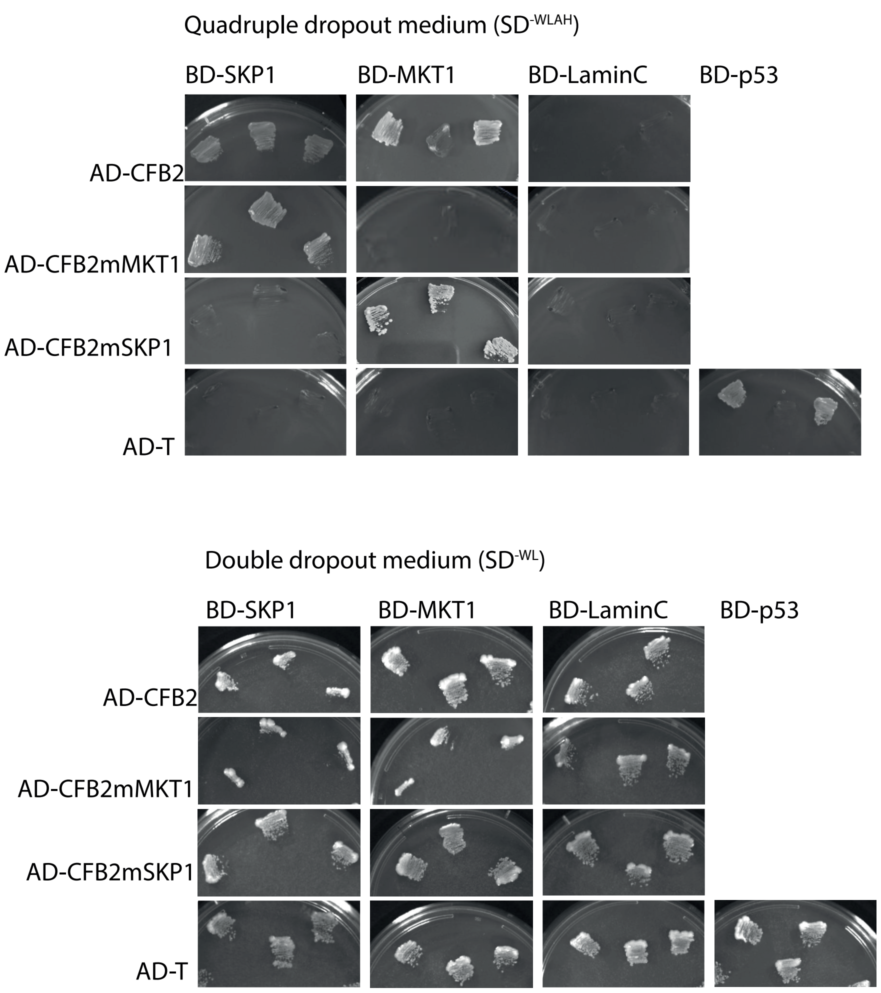


B.

| | pBD SKP1 | pBD MKT1 | pBD LaminC | pBD p53 |
|---|---|---|---|---|
| **pAD CFB2 wt** | 1 | 2 (*) | 3 (C-) | |
| **pAD CFB2 mMKT1** | 4 | 5 | 6 (C-) | |
| **pAD CFB2 mSKP1** | 7 | 8 | 9 (C-) | |
| **pAD T** | 10 (C-) | 11 (C-) | 12 (C-) | 13(C+) |

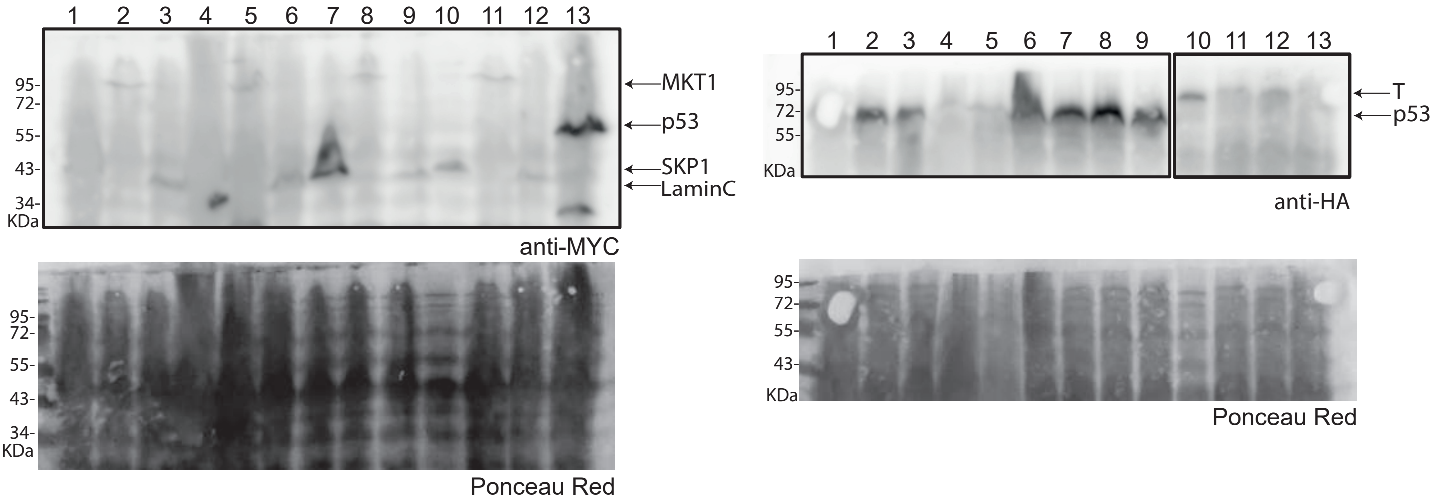

Fig. 4

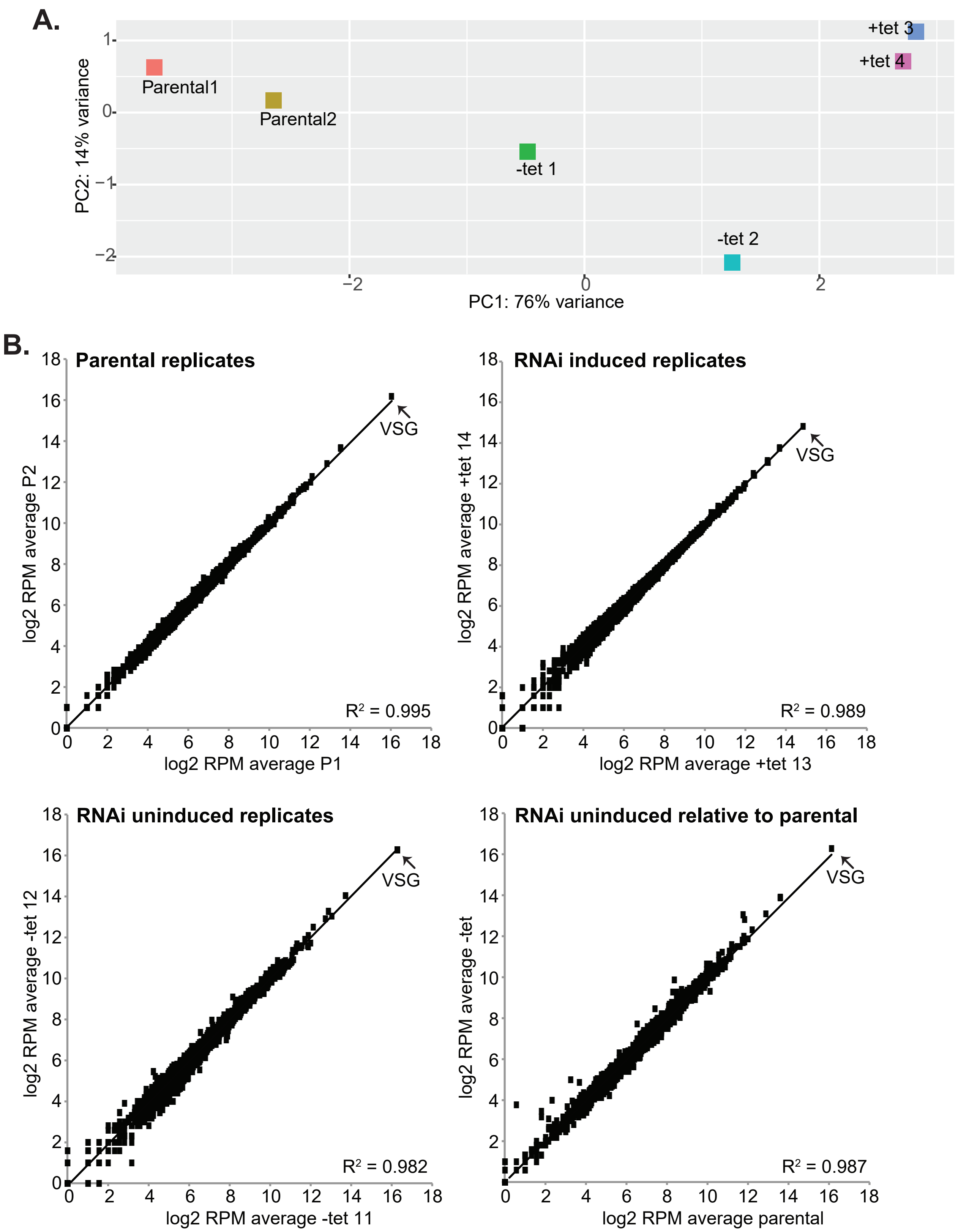
A.
PC2: 14% variance
PC1: 76% variance
Parental1
Parental2
-tet 1
-tet 2
+tet 3
+tet 4
B.
Parental replicates
log2 RPM average P2
log2 RPM average P1
VSG
$R^2$ = 0.995
RNAi induced replicates
log2 RPM average +tet 14
log2 RPM average +tet 13
VSG
$R^2$ = 0.989
RNAi uninduced replicates
log2 RPM average -tet 12
log2 RPM average -tet 11
VSG
$R^2$ = 0.982
RNAi uninduced relative to parental
log2 RPM average -tet
log2 RPM average parental
VSG
$R^2$ = 0.987

**Fig. 5**

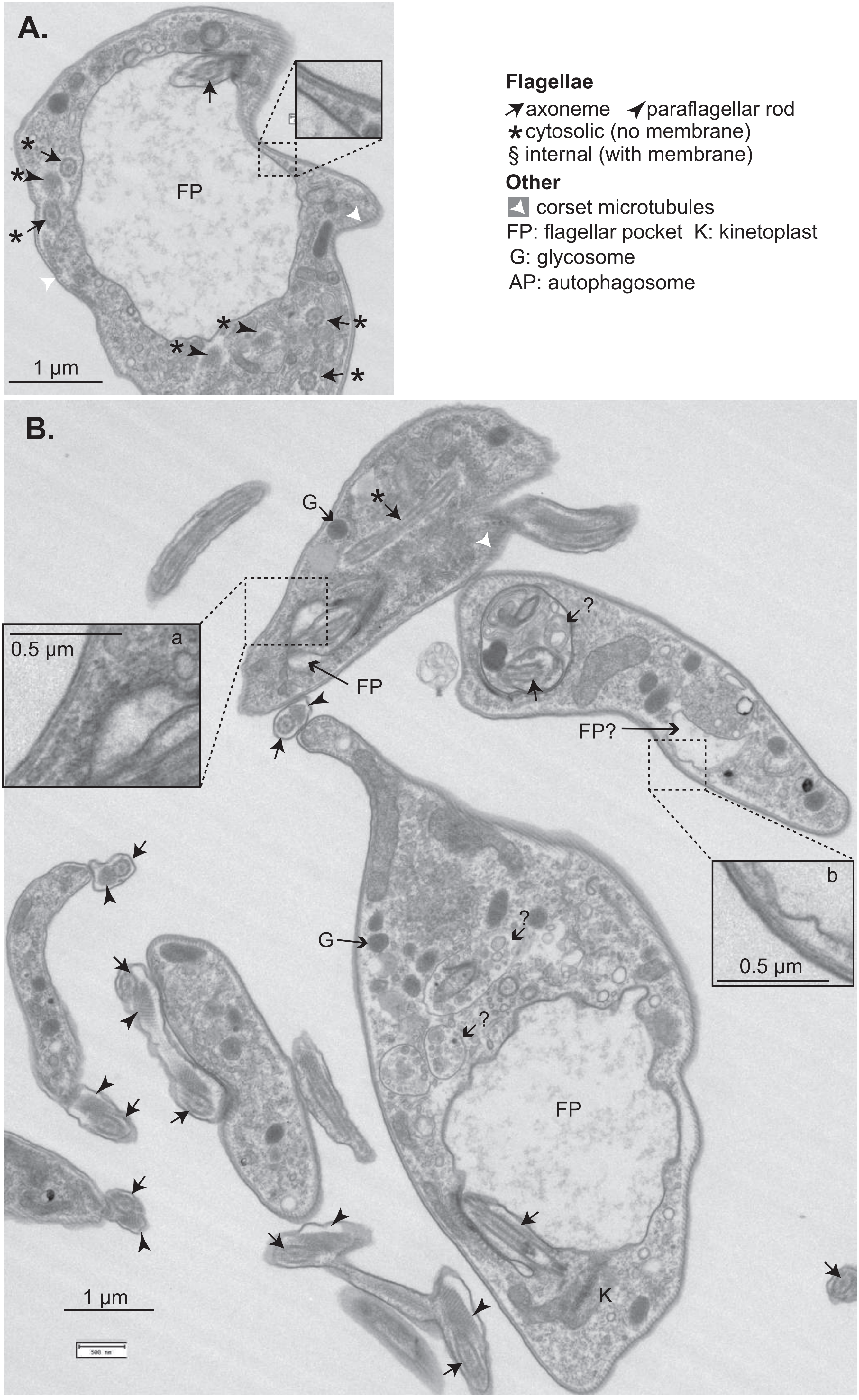
A.
FP
1 μm
Flagellae
axoneme
paraflagellar rod
cytosolic (no membrane)
§ internal (with membrane)
Other
corset microtubules
FP: flagellar pocket K: kinetoplast
G: glycosome
AP: autophagosome
B.
G
FP
a
0.5 μm
FP?
b
0.5 μm
G
FP
K
1 μm

**Fig. 6**

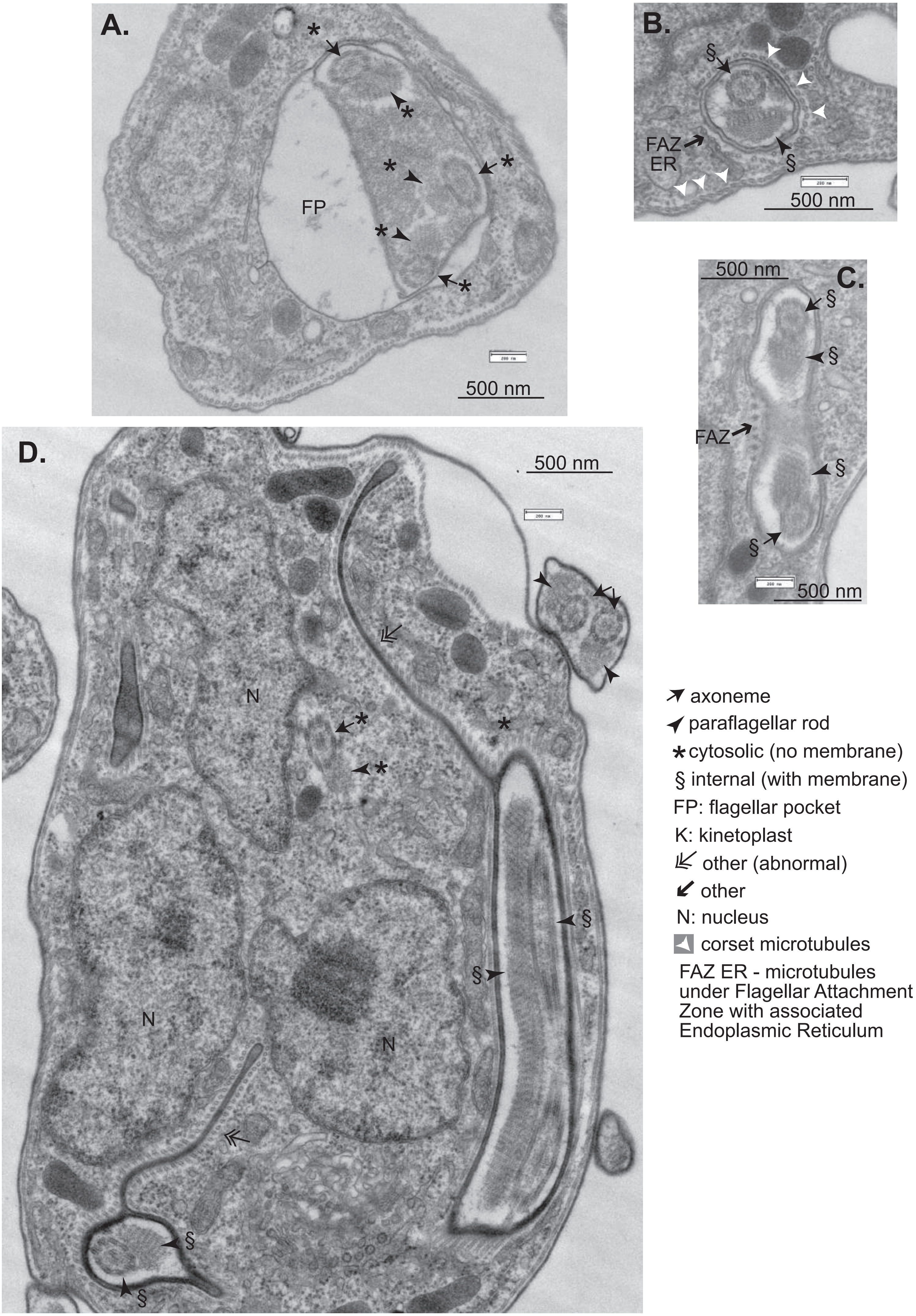
A.
FP
500 nm
B.
FAZ ER
500 nm
C.
500 nm
FAZ
500 nm
D.
500 nm
N
N
N
axoneme
paraflagellar rod
cytosolic (no membrane)
§ internal (with membrane)
FP: flagellar pocket
K: kinetoplast
other (abnormal)
other
N: nucleus
corset microtubules
FAZ ER - microtubules under Flagellar Attachment Zone with associated Endoplasmic Reticulum

Fig. 7

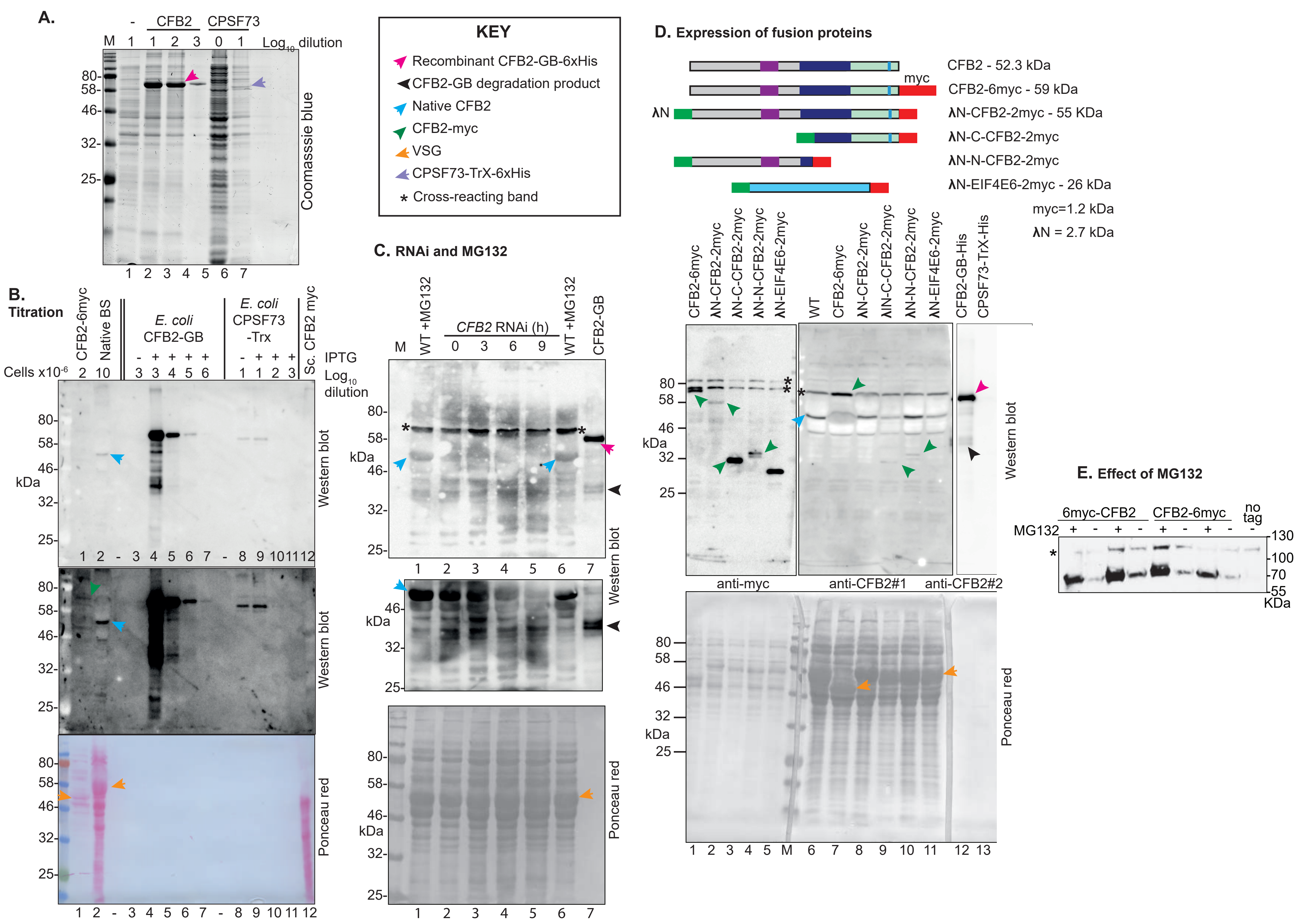

A.
CFB2
CPSF73
Log10 dilution
Coomasssie blue
KEY
Recombinant CFB2-GB-6xHis
CFB2-GB degradation product
Native CFB2
CFB2-myc
VSG
CPSF73-TrX-6xHis
* Cross-reacting band
D. Expression of fusion proteins
CFB2 - 52.3 kDa
CFB2-6myc - 59 kDa
λN-CFB2-2myc - 55 KDa
λN-C-CFB2-2myc
λN-N-CFB2-2myc
λN-EIF4E6-2myc - 26 kDa
myc=1.2 kDa
λN = 2.7 kDa
B. Titration
CFB2-6myc
Native BS
E. coli CFB2-GB
E. coli CPSF73 -Trx
Sc. CFB2 myc
IPTG
Cells x10-6
Log10 dilution
Western blot
Ponceau red
C. RNAi and MG132
WT +MG132
CFB2 RNAi (h)
CFB2-GB
kDa
WT
anti-myc
anti-CFB2#1
anti-CFB2#2
CFB2-GB-His
CPSF73-TrX-His
E. Effect of MG132
6myc-CFB2
CFB2-6myc
no tag
MG132
KDa

**Fig. 8**

**A.**

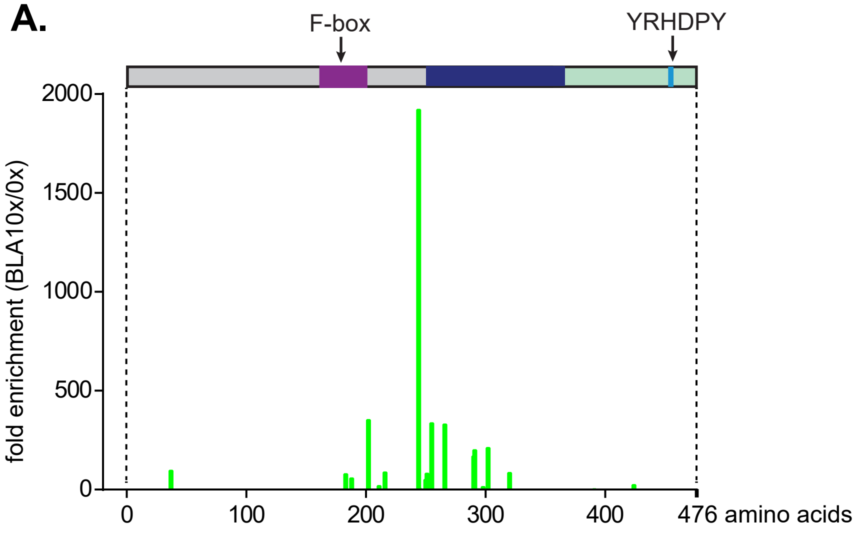


**B.**

```
Pcon   KLIRLFFGQQRRDSSPRALRRLLDYLVPDVTIPHVEAHTNKQSGRGKGCAWVLVTNEHDK
Tco70  KPIRLFFGQQRRDAHPSALRRLLRFIAPSLPIMHMEAHVNPVTGRGKGCVWATVPSLAEA
Tv67   KPIRLFFGQQRRDAHPSALRRLLRFIAPSLPIMHMEAHVNPVTGRGKGCVWATVPSLAEA
Tv45   KPIRLFFGQQRRDAHASALRRLLRFIVPSLPIMHMEAHTNSRNGRGKGCAWVIVPSLAEA
Tco60  KPIRLYFGNQRRDVYPSALRRLLRFIAPDLLITHMEGHANSGTGRGKGCAWVVVPSVAEA
TbCFB1 KPIRLFFGQQRRDPHPSALRRLLRFIAPDLVIAHMEAHTNPVNGRGKGCAWVFALSQLDA
TbCFB2 KPIRLFFGQQRRDVYPSALRRLLRFIAPDLVITHMEAHVNEATGRGKGCAWVIVPSVLEA
Tcr    KNIRLFFGQQRRDPHPASLRHLLNFLVPEISIPHMESHTNALNNRGKGCTWVFVTSQDDV
Tra    KTIRLFFGQQRRDNHPSSLRHLLNFLVPDVLIPQMEAHTNALNGRGKGCTWVFVTSQEDA
Tth    KNIRLFFGQQRRDNHPSSLRHLLNFLHPDMVIPHMEAHTNSLSGRGKGCSWVFVTSREDA
Tgr    KTIRLFFGQQRRDNHPSSLRHLLNFLVPDMAIPHMEAHTNPLNGRGKGCSWVFVTSQSDA
       * ***:**:****    :**:** :: *.: * ::*.*.*  ..***** *. . .  :

Pcon   ERLCRLNRRLFIDTAGP (x)42 AGERILLAPPREAALMALAEYAIRAGACTQRPNHLPRQLVVV
Tco70  KRLMALNKRLFLDVNSD------GEEVYMFAPPSAVE--WLRGHAELAAASTSRPSHLPRQPMIV
Tv67   KRLMALNKRLFLDVNSD------GEEVYMFAPPSAVE--WLRGHAELAAASTSRPSHLPRQPMIV
Tv45   KRLMALNRRLFLDVNSD------GEEVYMFAPPSAKE--WLHEHAELAAASTSRPNHLPRQPMVV
Tco60  KRLLLLSGRIFLDVNSD------GEEVYMFGPLTCRK--WLHNYAEHVTEVTPRQNHLPRQPMVV
TbCFB1 ERLLRLSGRIFLDINSN------GEEVYLFAPPNCRE--WLNEHAECAVASAVRPSHLPRQPMVV
TbCFB2 KRLLRLSGRIFLDINSN------GEEVYLFAPPNCRE--WLSEYADYVVSSTTRASHLPYLPMVV
Tcr    KRLMRFNRLIFLDLNAD------GREVYMVAPPRSRM--WIQEYAENIARWSTRPIYLPRQPMVI
Tra    KRLMQFNKLIFLDVNAD------GKEVYMVAPPRSRA--WLQDRAESVGRWATRPMCLPRQPMVI
Tth    SRLMAFNKLLFLDMREN------GEEVYLLAPPRSKD--WLQEYSQEVSTASYRSSNLPRQPMVV
Tgr    KRLMGFNKLLFLDVNER------GEEVYLLAPPCSKA--WLQKHAQDVGGRPFRPNNLPRQAVVV
       .**  :.  :*:*          . *  :..*        :   :        *   **   :::
```

Fig. 9

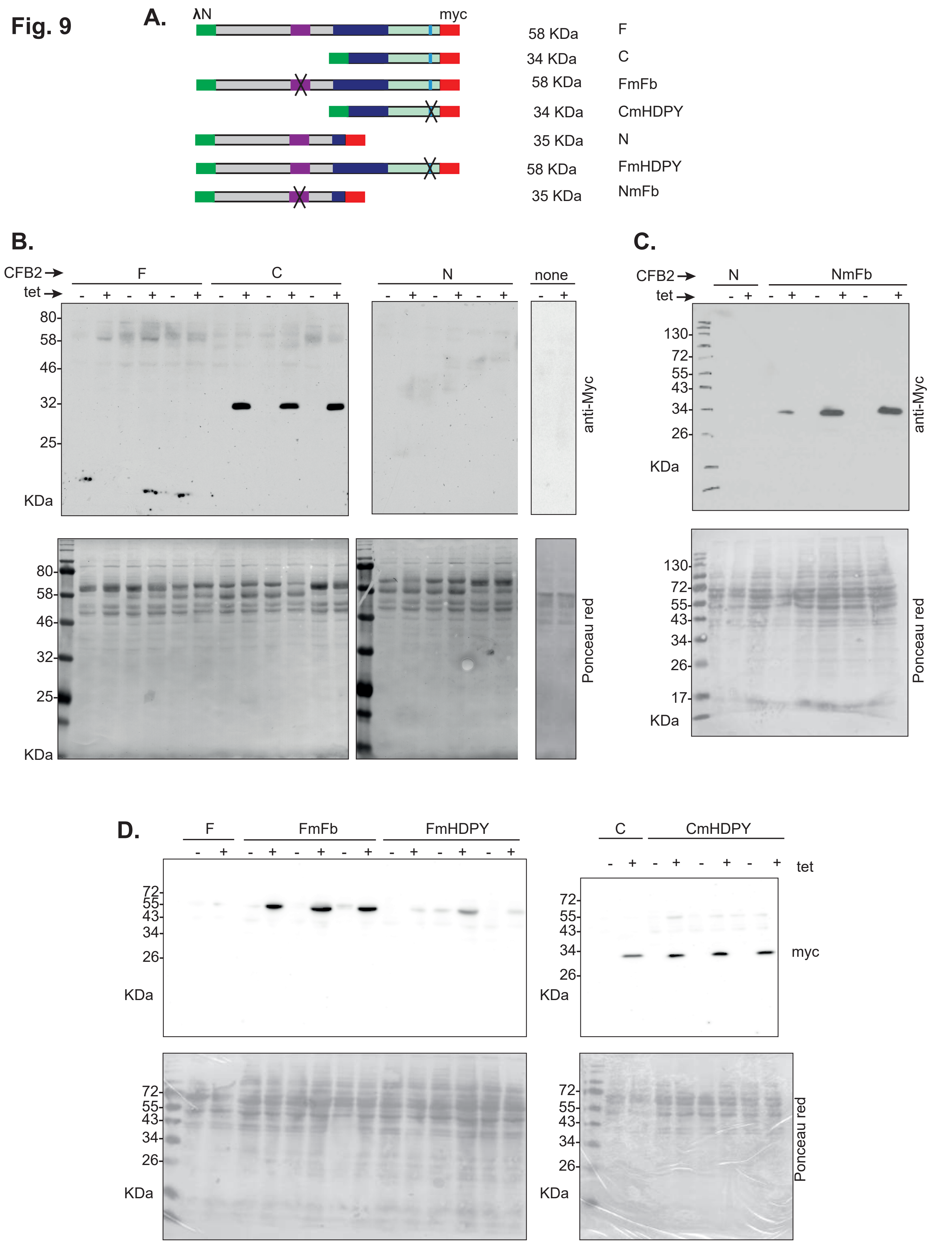
A.
λN
myc
58 KDa F
34 KDa C
58 KDa FmFb
34 KDa CmHDPY
35 KDa N
58 KDa FmHDPY
35 KDa NmFb
B.
CFB2
tet
F
C
N
none
80
58
46
32
25
KDa
anti-Myc
Ponceau red
C.
N
NmFb
130
72
55
43
34
26
17
D.
F
FmFb
FmHDPY
C
CmHDPY
tet
myc

### Fig. 10

```
VSG4 5' AAUUUUCCCCCUC--UUUUUC-----UUAAAAA--UUCUUGCUACUUGAAAAC-UCCUGAUAUAUUUUAACAC-AAAA 3'
VSG2 5' UAAUUCCCCCCUCAAUUUCCCCCUCCUUUUAAAUUUUCCUGCUACUUGAAAACUUUUUGAUAUAUUUUAACACCAAAA 3'
```

```
IlTat1.3        --------UAAUCCCUUUCCCC-UC-U--UUUCUUCC-AUGCUAAAAUUCU
IlTat1.24       UAA--AACAAAUAUAUCCCCCC-UC-A--AUUUUCCUUA---A--------
IlTat1.25       UAAAA--CAAAUAUAUCCCCCC-UC-A--AUUUUCCUUA---A--------
IlTat1.23       ----------UAAUUUUUCACC-UC-U----UUUUCUU----------CCU
VSG1125.4959    -------------AUUCCCCCC-UC-U--UUUUUUCUUA--AAGAAAAUUU
EATRO1125.2523  ---------------UUCCCCC-UC-U----UUUUCUUGCUAAAAUUUCCU
VSG1125.211     -------------UUUUCCCCC-UC-U--U----------------UUUCU
EATRO1125.313   --------AACAAUUUCCCCCC-UC-A--AUUUUUCUUU-UAAAAAAUUCU
EATRO1125.4710  ---AA--CAA--AUUUUCCUCC-UC-AA--UUUU-----------------
EATRO1125.1654  ---AA--CAA--AUUUUUCCCC-CC-UA--UUUU-----------------
VSG118          --UAA--CAA--AUUUUUCCCC-UC-AA--UUUU-----------------
MiTat1.5        UAAAA--CAA--AUUUUUCCCC-UC-AA--UUUU-----------------
AnTat11.17      -UAAA--UAA--UUUUCCCCCC-UC-----UUUUUCUUGCUGAA----AUU
VSG1125.4862    --------UU--UUUUUCCCCC-UC-A--ACUUUU--UAAGAAA----CUU
EATRO1125.2564  ---------A--AUUUUCCCCC-UC-UCAAUUUUUCUUUCUGAA----AUU
MVSG5           ---------U--UUUUUCCCCC-UC-UU--UU------UCU----------
BUGC18          -UAAA--CAA--AAUUUCCCCC-UC-AA--UU------UUU----------
VSG117=VSG4     --------UA--AUUUUCCCCC-UC-UU--UU------UCU----------
MiTat1.6        --------UA--AUUUUCCCCC-UC-UU--UU------UCU----------
VSG222a#        ---------UAAUCCCUCCCCCCCCCCCGGA-AAAUUU--------UCCU
VSG222b#        ---------UAAUCCCUCCCCCCUCCUUUUA-AAAUUU--------UCCU
AnTat1.8        ----------UAAUUAGCCCCCC----------UCUUUCUAAAAAUUUUC
EATRO1125.514   ----------UUUUUA-ACCCC------------------------UCUU
EATRO1125.385   -----------------CCCCC------------------------UCCG
IlTat1.1        --------UAA-UUUUUCCCUU---UAAAAUUU---UCC-------UCCU
MiTat1.1        ------------UAAUUCCCCC------------------------UCUU
EATRO1125.276   ---AA---CAA-UUUUUCCCCC-UC--AAAUUC-CCCUC-------UCUU
EATRO1125.421   --------CAA-A-UUUCCCCC---CCAAAUUC--CCCC-------UCUU
IlTat1.22       UAAAA--CAAA-AUUUUCCCCC----AAAUUCC--CCAC-------UCUU
MiTat1.2=VSG2   ---------UA-A-UUCCCCCC---UCAAUUUC--CCCC-------UCCU
IlTat1.21       ---------UA-AUUUUCCCAC---UCAAAUUC--CCUC-------UCCC
EATRO1125.84    -------------UUUUCCCAC---UCAAAUUC--CCUC-------UCCC
```

```
IlTat1.3        UGCUAAA--AAAUUUUGCUACUUGAAA--ACUU-----UCUGAUAUAUUUUAACACCU-
IlTat1.24       -----AA--AAUUCUUGCUACUUGAAA--ACUU-----UCUGAUAUAUUUUAACACGUAAGUUAAAAAAAAAAAA
IlTat1.25       -----AA--AAUUCUUGCUACUUGAA---ACUU-----UCUGAUAUAUUUUAACACGUAAGUUACCCGAAAAAAA
IlTat1.23       UGCUAAA--A-AUAUUGCUACUUGAAA--ACUU-----UCUGAUAUAUUUUAACAC---
VSG1125.4959    UGCUAAA--AUUUUUUGCUACUUGAAAAAACUU-----UCUGAUAUAUUUUAACACGUAAGUUACCCGAAAAAAA
EATRO1125.2523  UGCUAU---AAUUUUUGCUACUUGAAA-A-CUU-----UCUGAUAUAUUUUAACACGCAAGUUACCCGAG
VSG1125.211     UGCUAAA--AAUUCUUGCUACUUGAAA-AAUUU-----UCUGAUAUAUUUUAACACGCAAGUUACCCGAAAAAAA
EATRO1125.313   UGCUAAA--AAUUCUUGCUAUUUGAAA-A-CUU-----UCUGAUAUAUUUUAACACGCAAGUUACCCGAAAAAAA
EATRO1125.4710  --UAAAAA-GAAUUUUCCUACUUGAAAA-ACUU-----UCUGAUAUAUUUUAACACGCG
EATRO1125.1654  --UAAAAA--AUUCUUGCUAUUUGAAAA-ACUU-----UCUGAUAUAUUUUAACACCUAAAACCAGCCGAAA
VSG118          -----AAA--AUUUUUGCUACUUGAAAA-ACUU-----UCUGAUAUAUUUUAACACGUAAGUU
MiTat1.5        -----AAA--AUUUUUGCUACUUGAAAA-ACUU-----UCUGAUAUAUUUUAACAC---
AnTat11.17      UACUA-AA-AUUUUUUGCUACUUG-AAA-ACUU-----UCUGAUAUAUUUUAACACCU-
VSG1125.4862    UGCUAAAA-AUUUCUUGCUACUUG-AAA-ACUU-----UCUGAUAUAUUUUAACACCUAAAAAAA
EATRO1125.2564  UGCUAAAA-AAUUCUUGCUACUUGAAAA-ACUU-----UCUGAUAUAUUUUAACACCUAAAAAAA
MVSG5           --U-AAAG-AAUUUUUGCUACUUUAAAA-ACU------UCUGAUAUAUUUUAACACCUAAAACCAACCGAGC
BUGC18          --AAAAAG-AAUUUUUGCUAUUUG-AAA-ACU------UCUGAUAUAUUUUAACACCCC
VSG117=VSG4     ----UAAA-AAUUCUUGCUACUUG-AAA-ACU------CCUGAUAUAUUUUAACA----
MiTat1.6        ----UAAA-AAUUCUUGCUACUUG-AAA-ACU------CCUGAUAUAUUUUAACAC---
VSG222a#        UGCUA--ACAUUUUUUGCUACUUGAAAA-ACUU-----UCUG-UAUAUUUUAACGAAUUC
VSG222b#        UGCUA--ACAUUUUUUGCUACUUGAAAA-ACUU-----UCUGAUAUAUUUUAACGAAUUC
AnTat1.8        CCCCUGCUAAAAUUUUGCUACU-----------------CUGAUAUAUUUUAACACCUAAAAAAUUCCCCG
EATRO1125.514   UUUAAA-AUUUUCCUUGCUA----AAAA-UUUUUGCUAUUUGAUAUAUUUUAACACGUAA
EATRO1125.385   UAAAGA---A--CUUUGCUACUUGAAAA-CCUU-----UCUGAUAUAUUUUAACACCUAAAAAAUCAGCCGAAAAA
IlTat1.1        UUAAAAACAAAUUUUUUCUACUUGAAAA-CUUCUGAUAUUUGAUAUAUUUUAACACCUUUU
MiTat1.1        UUUAAACGAA--AUUUGCUA-UU------------------UGAUAUAUUUUAACAC---
EATRO1125.276   UUCAAAAAUCUUCCUUGCUACUUGAAAC-CUU------UCUGAUAUAUUUUAACACCUUU
EATRO1125.421   UUUAAA-AAA--CUUUGCUACUUGAAAA-ACU------UUUGAUAUAUUUUAACACGUAAUUUCUGCCGAAAAA
IlTat1.22       UUUAAA-AUUUUCCUUGCUACUUGAAAA-CUU------UCUGAUAUAUUUUAACACCCAAAAAAA
MiTat1.2=VSG2   UUUAAA-UUU--UCCUGCUACUUGAAAA-CUU------UUUGAUAUAUUUUAACACCAAAAAAA
IlTat1.21       UUUAAA-AAC--U-UUGCUACUUGAAAA-CUU------UUUGAUAUAUUUUAACACCUGAAAAAAA
EATRO1125.84    UUUAAA-AAC--U-UUGCUACUUGAAAA-CUU------UUUGAUAUAUUUUAACACCUGAA
                              *****                   ** ***********
                             "8-mer"                   "16-mer"
```

#a: Sequence from cloned cDNA; b: sequence from genomic sequencing (Tb427_2018)

Fig. 11

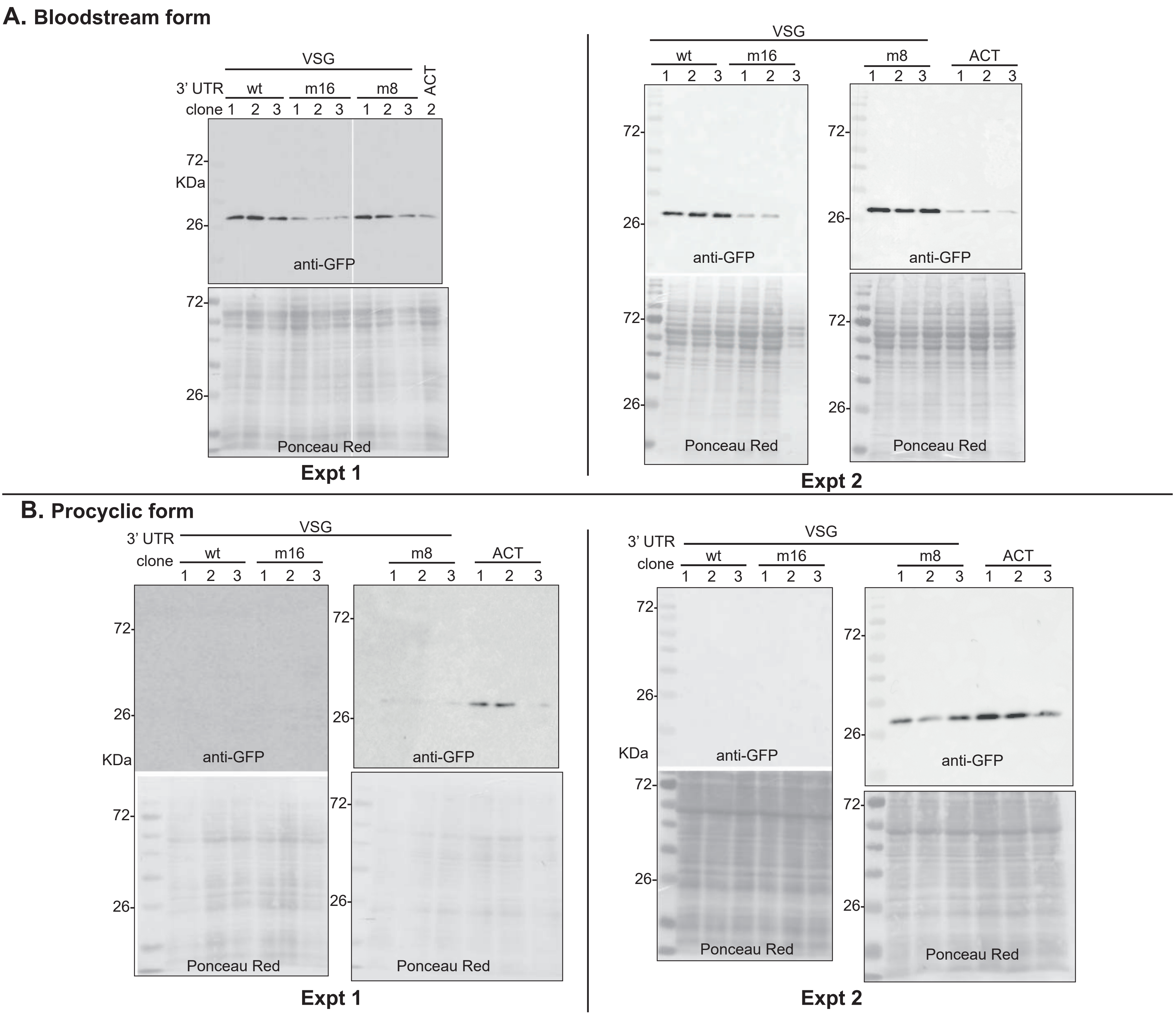
A. Bloodstream form
VSG
3' UTR
wt
m16
m8
ACT
clone
1 2 3 1 2 3 1 2 3 2
72-
KDa
26-
anti-GFP
Ponceau Red
Expt 1
Expt 2
B. Procyclic form
anti-GFP
Ponceau Red
Expt 1
Expt 2

Fig. 12

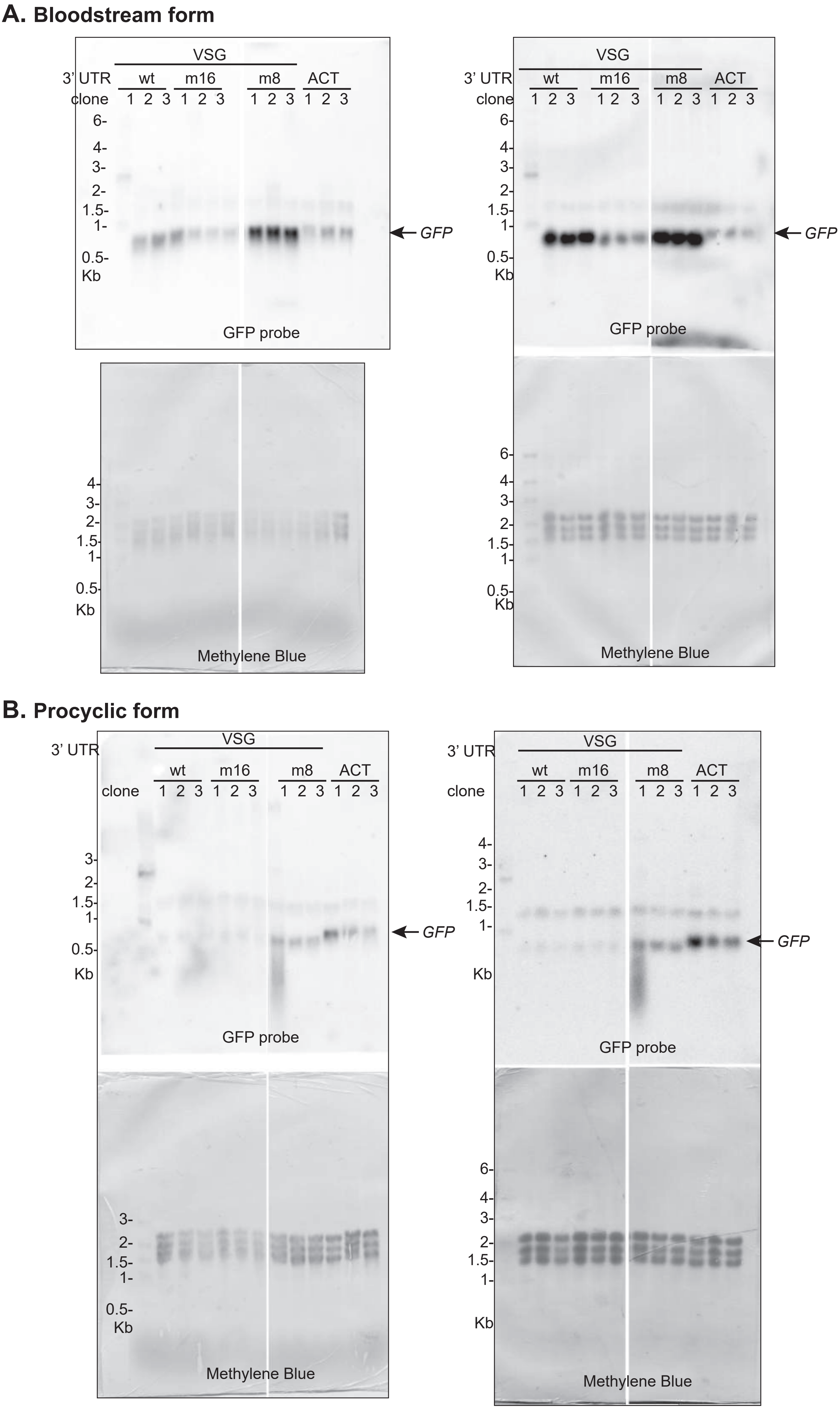

A. Bloodstream form
VSG
3' UTR
wt
m16
m8
ACT
clone
1 2 3 1 2 3 1 2 3 1 2 3
6-
4-
3-
2-
1.5-
1-
0.5-
Kb
GFP
GFP probe
Methylene Blue
B. Procyclic form
VSG
3' UTR
wt
m16
m8
ACT
clone
1 2 3 1 2 3 1 2 3 1 2 3
GFP
GFP probe
Methylene Blue

Fig. 13

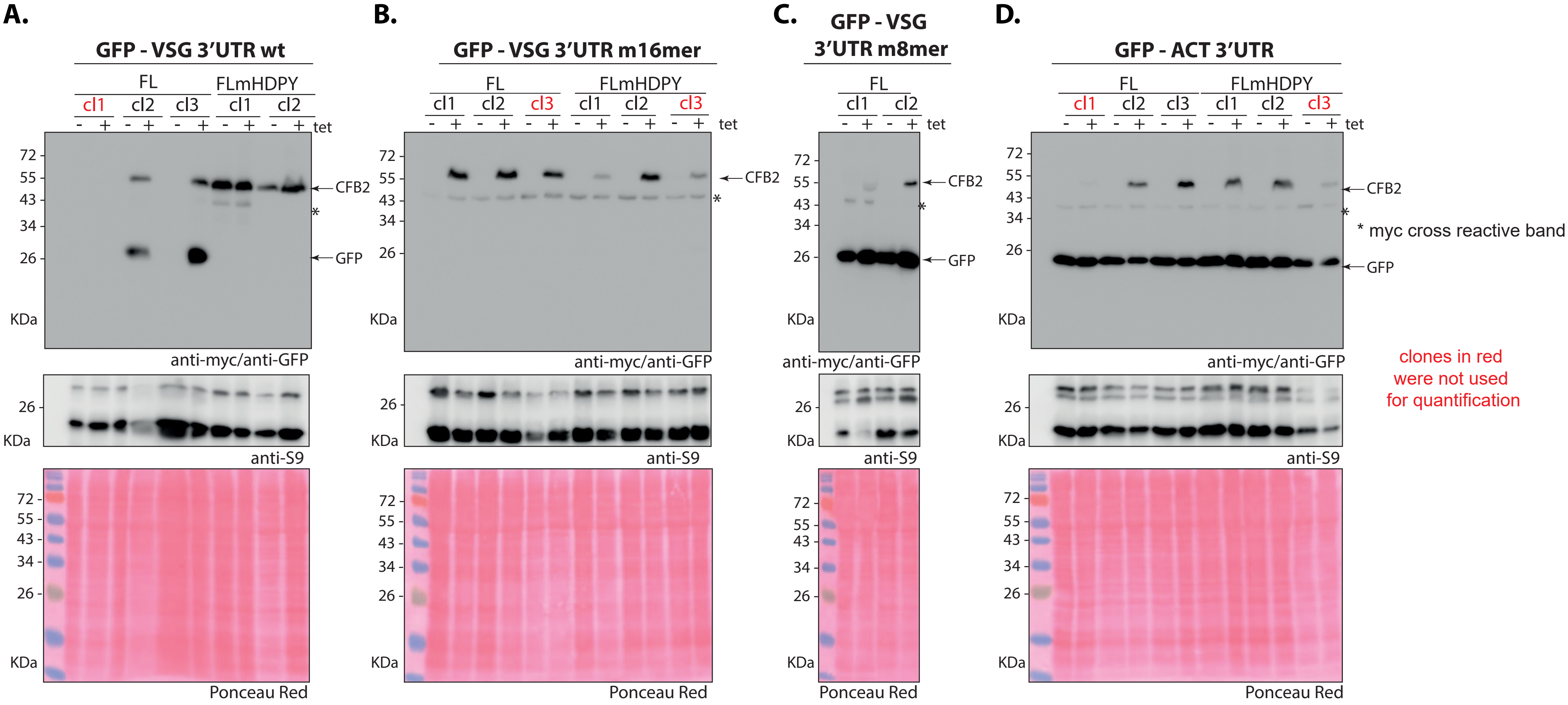
A.
GFP - VSG 3'UTR wt
FL
FLmHDPY
cl1
cl2
cl3
cl1
cl2
- + - + - + - + - + tet
72
55
43
34
26
KDa
CFB2
*
GFP
anti-myc/anti-GFP
26
KDa
anti-S9
72
55
43
34
26
KDa
Ponceau Red
B.
GFP - VSG 3'UTR m16mer
FL
FLmHDPY
cl1
cl2
cl3
cl1
cl2
cl3
- + - + - + - + - + - + tet
72
55
43
34
26
KDa
CFB2
*
anti-myc/anti-GFP
26
KDa
anti-S9
72
55
43
34
26
KDa
Ponceau Red
C.
GFP - VSG 3'UTR m8mer
FL
cl1
cl2
- + - + tet
72
55
43
34
26
KDa
CFB2
*
GFP
anti-myc/anti-GFP
26
KDa
anti-S9
72
55
43
34
26
KDa
Ponceau Red
D.
GFP - ACT 3'UTR
FL
FLmHDPY
cl1
cl2
cl3
cl1
cl2
cl3
- + - + - + - + - + - + tet
72
55
43
34
26
KDa
CFB2
*
* myc cross reactive band
GFP
anti-myc/anti-GFP
clones in red
were not used
for quantification
26
KDa
anti-S9
72
55
43
34
26
KDa
Ponceau Red
